# Structural Brain Network Alterations in Relation to Treatment and Illness Severity in Bipolar Disorder

**DOI:** 10.64898/2026.03.28.714565

**Authors:** Leila Nabulsi, Melody J.Y. Kang, Neda Jahanshad, Genevieve McPhilemy, Fiona M. Martyn, Bartholomeus Haarman, Colm McDonald, Brian Hallahan, Stefani O’Donoghue, Dan J. Stein, Fleur M. Howells, Freda Scheffler, Henk S. Temmingh, David C. Glahn, Amanda Rodrigue, Edith Pomarol-Clotet, Eduard Vieta, Joaquim Radua, Raymond Salvador, Andriana Karuk, Erick J. Canales-Rodríguez, Josselin Houenou, Pauline Favre, Mircea Polosan, Arnaud Pouchon, Paolo Brambilla, Marcella Bellani, Philip B. Mitchell, Gloria Roberts, Udo Dannlowski, Tiana Borgers, Susanne Meinert, Kira Flinkenflügel, Jonathan Repple, Elisabeth J. Leehr, Dominik Grotegerd, Tim Hahn, Michèle Wessa, Mary L. Phillips, Lea Teutenberg, Tilo Kircher, Benjamin Straube, Olaf Steinstraeter, Frederike Stein, Florian Thomas-Odenthal, Nina Alexander, Paula L Usemann, Andreas Jansen, Michael Berk, Orwa Dandash, Nadine Parker, Chao Suo, Sophia I. Thomopoulos, Paul M. Thompson, Ole A. Andreassen, Christopher R. K. Ching, Dara M. Cannon, ENIGMA Bipolar Disorder Working Group

## Abstract

**Background:** Large-scale T1-weighted MRI studies have established grey-matter abnormalities in bipolar disorder (BD), with our group contributing to consensus findings. However, structural connectivity, particularly within emotion- and reward-related circuits, remains poorly understood. Diffusion-weighted MRI (dMRI) enables investigation of white-matter pathways, yet prior work is constrained by small samples, methodological heterogeneity, and unclear medication effects. We conducted the largest dMRI network analysis in BD, relating symptom burden and polypharmacy to tractography-derived connectivity and graph-theoretic metrics.

**Methods:** Cross-sectional structural and diffusion MRI scans from 449 individuals with BD (35.7±12.6 years) and 510 controls (33.3±12.6 years), aged 18–65, were analyzed across 16 ENIGMA-BD sites. Standardized segmentation/parcellation and constrained spherical deconvolution tractography generated individual structural connectivity matrices. Graph-theoretic metrics of global and subnetwork organization were related to symptom severity and medications.

**Results:** BD showed widespread network alterations (lower density and efficiency, longer path length, and higher betweenness centrality), altered microstructural organization in a limbic–basal ganglia circuit, and abnormal streamline counts in a default-mode/salience/fronto-limbic–basal ganglia network. Longer illness duration, later onset, and psychosis history were associated with greater abnormalities in network architecture, whereas more manic episodes were associated with greater fronto-limbic connectivity. Antidepressant (particularly SSRI), anticonvulsant, and antipsychotic use related to poorer global and fronto-limbic connectivity; no clear lithium effects emerged.

**Conclusions:** As the largest structural connectivity study in BD, we reveal widespread disruption in reward and emotion-regulation networks influenced by illness severity and medication use. Results show that multisite harmonization is feasible and highlight ENIGMA-BD as a scalable framework for identifying reproducible neurobiological markers.

## Introduction

Bipolar disorder (BD) is a chronic, recurrent mood disorder characterized by depression and (hypo)mania, affecting roughly one in 25 adults in the United States and contributing substantially to global disability (Vos et al., 2017). Although cognitive and affective impairments are well-documented, treatment response remains highly variable: only ∼30% of patients respond robustly to lithium–the standard first-line agent (Malhi et al., 2013)–and many continue to experience persistent symptoms, functional impairment, or treatment resistance (McIntyre & Calabrese, 2019; Levenberg & Cordner, 2022). BD also carries one of the highest suicide rates among psychiatric disorders (Chesney et al., 2014). Mapping neural circuit dysfunction is therefore critical for understanding illness burden, predicting treatment response, and advancing biologically grounded, personalized strategies.

BD is characterized by dysregulation within fronto-limbic circuits and disrupted interactions with networks supporting fear, anxiety, and cognitive control (O’Donoghue et al., 2017; Ching et al., 2020). Diffusion-weighted MRI studies show that white matter (WM) abnormalities extend beyond anterior fronto-limbic pathways to major association and projection fibers—including the superior longitudinal (arcuate), inferior longitudinal, inferior fronto-occipital fasciculi, posterior thalamic radiations, and internal capsule (Favre et al., 2019; Nadine et al., 2025). Robust effects are also observed in the corpus callosum and cingulum, where severity relates to age of onset and illness duration (Favre et al., 2019), although the contribution of callosal transfer to mood switching and emotional regulation remains unclear (Wang et al., 2008; Linke et al., 2013). Altered connectivity in posterior and cerebellar pathways may reflect compensation for disrupted frontal circuits (Nabulsi et al., 2023). WM disruptions are particularly pronounced in psychotic BD presentations (Sarrazin et al., 2014) and may contribute to impaired cognitive control and affective instability.

Network neuroscience reframes brain function from regional localization to connectivity-based models, emphasizing distributed interactions among neural systems. Graph theory provides a mathematical framework for this approach, representing regions as nodes and white-matter pathways as edges, enabling quantification of network organization and integration (Sporns, 2013). Although diffusion and structural imaging studies report focal WM abnormalities in BD (Perry et al., 2018), prior graph-theoretic work has relied on small samples with heterogeneous pipelines, leading to inconsistent network findings. Variability in acquisition, segmentation, parcellation, and connectivity metrics, as well as differences in illness duration, severity, and medication exposure, further contributes to these discrepancies. Large-scale, harmonized network analyses that control for site effects are needed to generate more reproducible estimates of BD circuitry, clarify illness-related variation, and determine how common treatments shape network architecture.

Most patients with BD require the concurrent use of multiple medications (polypharmacy) to control their symptoms. Large multi-site studies in individuals with BD have reported associations between psychotropic medications and structural brain features. Lithium use has been linked to greater cortical thickness in parietal and frontal regions, while anticonvulsants and antipsychotics have been associated with patterns of thinner cortex and lower surface area in occipital and frontal regions (Hibar et al., 2016; 2017). In the first diffusion-weighted MRI (dMRI study from ENIGMA-BD including 3,033 individuals across 26 sites, Favre et al. (2019) found associations between common medications and WM microstructure using diffusion tensor imaging (DTI). Lithium was associated with patterns of higher fractional anisotropy (FA) and lower mean diffusivity (MD), while antipsychotics and anticonvulsants were linked to lower FA. Similar analyses using advanced WM connectivity features such as graph theory are needed to better understand the relationship between network dysfunction, treatment and illness severity in BD.

The Enhancing Neuro Imaging Genetics through Meta-Analysis Bipolar Disorder Working Group (ENIGMA-BD) is the largest global consortium for harmonized neuroimaging and clinical data in BD, enabling standardized processing and improved replicability of brain signatures (Thompson et al., 2020; Ching et al., 2020). Using a standardized pipeline across 16 datasets (Figure S1), we assessed whole-brain and subnetwork structural connectivity in BD versus controls. We hypothesized that BD would show network-level disruptions: reduced clustering and efficiency, longer characteristic paths, and higher betweenness centrality, alongside alterations within limbic, basal ganglia, interhemispheric, and cerebellar projections (Favre et al., 2019; Nabulsi et al., 2019; 2023). We further predicted that greater symptom burden would relate to more pronounced abnormalities, as previously described (Favre et al., 2019), and that medication exposure, particularly lithium, anticonvulsants, and antipsychotics (especially when categorized by pharmacological mechanisms of action) would account for additional variance in WM network organization, consistent with prior ENIGMA findings (Favre et al., 2019; Hibar et al., 2016; 2017). Overall, our goal was to examine the relationships between network metrics, treatment responses, and symptoms, to advance the current understanding of BD-related structural networks and identify biologically grounded targets for interventions.

## Methods and Materials

### Subjects

Sixteen independent cohorts from ENIGMA-BD participated in the study, including 959 participants (449 individuals with BD and 510 controls; female/male: 51.1/48.9%). Demographics and clinical information are detailed in Table 1. Participants were between 18 and 65 years of age and fulfilled criteria for BD type I or II during clinical interview with a psychiatrist by DSM-IV-R criteria. Each cohort’s demographics and clinical information are detailed in Table S2. Individuals with BD type I (BD-I N=201, 83%) and type II (BD-II N=42, 17%) were included in the analysis. Both subtypes were combined for the main analysis, with follow-up tests conducted to examine differences between BD-I and BD-II. Analyses focused on adults (18-65), an age range commonly used in prior studies and best represented across sites; participants younger than 18 or older than 65, or with poor-quality MRI data after inspection were excluded from our analyses. Site-specific inclusion/exclusion criteria are detailed in Table S3. All participating sites obtained approval from their local institutional review boards and ethics committees, and all study participants provided written informed consent.

**Table 1.** Clinical and Sociodemographic Details of Participants. n/N = Sample/Total Sample Size; SD = Standard Deviation. *t* = t-test. *χ²*= chi-square.

|  | Controls<br>N = 509 | Bipolar Disorder<br>N = 450 | Statistical Comparison<br>Diagnostic Groups |
| --- | --- | --- | --- |
| Age, Mean (SD) | 33.25 (12.61) | 35.65 (12.59) | $t_{943} = 2.9, p = 0.003$ |
| Sex, n (%) | | | $\chi^2_1 = 6.9, p = 0.008$ |
| Male | 228 (45%) | 241 (54%) |  |
| Female | 281 (55%) | 209 (46%) |  |

### Image acquisition and processing

Contributing sites shared their raw T1-weighted and diffusion-weighted (dMRI) scans with the central processing team. Acquisition parameters for each site are detailed in Table S4. A probabilistic approach was employed to map subject-specific cortico-subcortical brain networks encompassing 34 cortical and 9 subcortical brain regions on both sides and including the cerebellum (FreeSurfer v5.3.0; Fischl 2012). This mapping was based on the Desikan-Killiany atlas (Desikan et al. 2006) applied to the T1-weighted image, covering a total of 86 regions (Figure 1). Diffusion MR images were processed using a deterministic constrained spherical deconvolution (CSD) algorithm was used to resolve crossing fibers within voxels (ExploreDTI v4.8.6; Tournier et al. 2007; Jeurissen et al. 2014), as described in Nabulsi et al. (2019) and detailed in *Supplementary Material*. 3D volumetric structural MR images were visually inspected before and after processing to ensure the accuracy of cortico-subcortical parcellation and segmentation, particularly examining gray and white matter boundaries. Parcellation/segmentation from T1-weighted images was evaluated using standard ENIGMA protocols (https://enigma.ini.usc.edu/protocols/imaging-protocols/; https://github.com/ENIGMA-git/ENIGMA-FreeSurfer-protocol). Diffusion MRI data was carefully inspected for artifacts, head motion, signal dropout, eddy-current-induced distortion and partial volume effects using in-house scripts. Structural connectivity matrices (86×86) were constructed and weighted by fractional anisotropy (FA), reflecting the average FA between two nodes, and by the number of streamlines (NOS), representing the number of reconstructed trajectories between nodes. These weights, generated during tractography, provide complementary information on microstructural organization (FA) and connection density (NOS). Binary (unweighted) matrices were also derived. Network topology was examined in both BD and control groups using FA/NOS-weighted and unweighted matrices.

**Figure 1.**
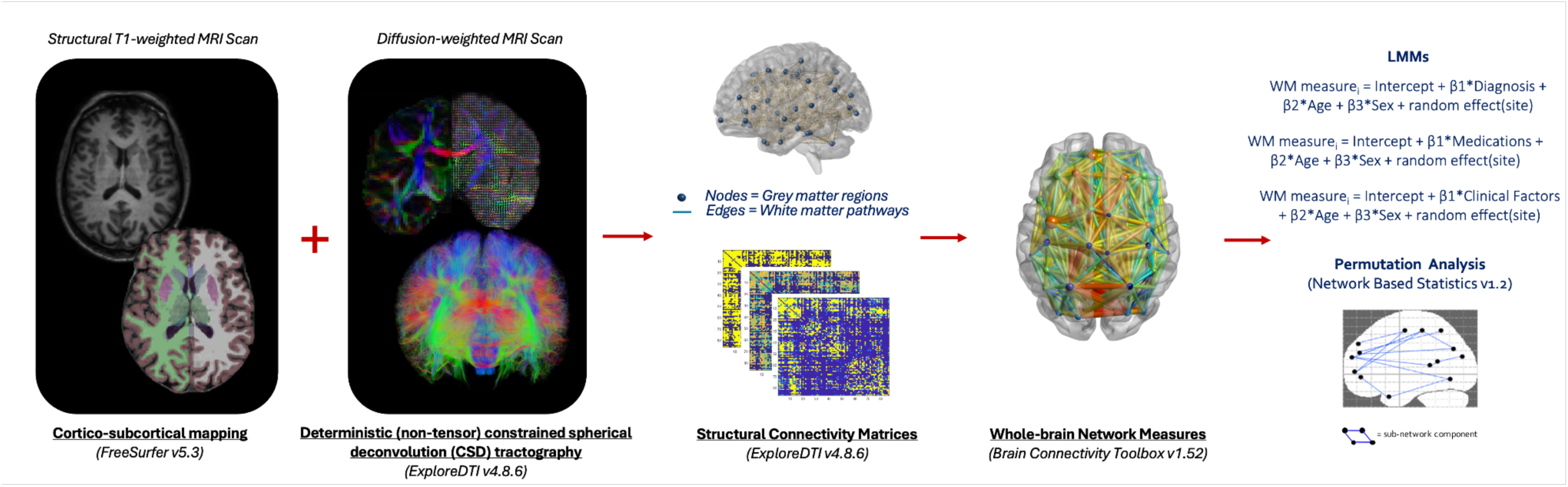
Human Connectome Reconstruction. This figure illustrates the standardized process of human connectome reconstruction and subsequent analyses. Proceeding from left to right: Brain Region Definition (node): The brain is divided into 34 cortical and 9 subcortical regions bilaterally using FreeSurfer v5.3.0, based on the Desikan-Killiany atlas. White Matter Trajectory Reconstruction (edge): White matter pathways are reconstructed using a deterministic non-tensor constrained spherical deconvolution algorithm, implemented via ExploreDTI v4.8.6. Brain Network Construction: The resulting brain network, composed of nodes and edges (human connectome) is represented as a structural connectivity matrix. Matrices are both unweighted and weighted by fractional anisotropy or the number of streamlines, generated using ExploreDTI v4.8.6. Global Network Measures: From the structural connectivity matrices, whole-brain network measures describing features of segregation and integration are calculated using the Brain Connectivity Toolbox v1.52). Linear mixed models (LMMs) are fitted to investigate structural connectivity differences and the influence of medications and clinical variables; a permutation analysis is performed to identify subnetwork-level differences between patients and controls, using NBS v1.2 (Zalesky et al. 2010).

### Whole-brain measures derived from the connectome

Global parameters summarizing whole-brain connectivity properties were extracted from both unweighted and weighted matrices (Figure 1). Measures of segregation, including global density and clustering coefficient were computed as the mean of the respective 86 regional estimates using functions from the Brain Connectivity Toolbox v1.52 (Rubinov and Sporns 2010). Measures of integration such as characteristic path length and efficiency were derived, as well as measures of influence and centrality, global degree/strength and betweenness (described in detail in Table S5).

### Statistical analysis of the structural connectome

We used linear mixed models (LMMs) to assess the relationship between whole-brain normally distributed measures and diagnosis, while adjusting for age, sex and site. In this model, diagnosis, age, and sex were included as fixed effects, and site was included as a random intercept to account for between-site variability (R v4.2.1) (Figure 1). We also tested for age-by-diagnosis and sex-by-diagnosis interactions for further evaluation of these effects. Multiple comparisons were accounted for using False Discovery Rate (FDR) correction (*p_FDR_*<.05) (Benjamini and Hochberg 1995), applied across all tests, consistent with prior work using similar graph-theory frameworks (Nabulsi et al., 2019; 2022). Network-Based Statistics (NBS v1.2; Zalesky et al., 2010) was used to perform mass univariate testing on FA- and NOS-weighted connectivity graphs (Figure 1), identifying subgraphs with significantly weaker or stronger connectivity while controlling for the family-wise error rate (FWER). A test statistic (F-test, adjusting for age, sex, and site) was computed to test for group connectivity strength differences (M=5000; *p*<.05). Connections were thresholded to obtain a set of suprathreshold *(t)* connections, namely only those connections that exceeded the set value. While the choice of primary threshold is user-defined, FWER correction through permutation testing ensures validity of results regardless of threshold choice (Zalesky et al., 2010).

### Clinical associations in bipolar disorder

Connectivity associations with clinical variables were tested in the BD group using linear mixed-effects models (pFDR < .05). For illness duration and age at onset, age was residualized to ensure effects were independent of age-related influences. Polypharmacy was examined using two medication frameworks: (1) conventional indication-based categories (lithium, antipsychotics, antidepressants, anticonvulsants; Table S1), and (2) mechanism-of-action groups based on Neuroscience-Based Nomenclature (NbN; Worley, 2017), which classifies medications by primary neurobiological targets rather than indication, using medication names or Anatomical Therapeutic Chemical (ATC) codes provided by each site (Supplement Table S6). Primary analyses focused on illness duration, age at onset, and lithium, anticonvulsant, and antipsychotic use; exploratory models included symptom-severity variables (e.g., psychosis history, episode counts) and NbN categories. All clinical models tested whether effects persisted when controlling for medication use, while medication models adjusted for concurrent medications (polypharmacy) and illness-course severity (the number of manic and depressive episodes), ensuring distinct contributions of clinical and pharmacological factors. Statistical model specifications are provided in Supplement Table S7.

## Results

### Participants’ clinical and demographic characteristics

Individuals with BD were predominantly euthymic (86%) at the time of scanning, significantly older than controls and comprised a higher proportion of females (55%) (Table 1). Clinical and sociodemographic details of BD are provided in Table S8.

### Whole-brain measures of network integration and segregation

BD and controls significantly differed in integration and segregation connectivity metrics (Figure 2; Table 2). Unweighted network topology in BD showed lower density, longer characteristic path length, lower global efficiency, and higher betweenness centrality (Cohen’s *d*=-.2–.2; *p_FDR_*=[.01-.04]). Similar disruptions were observed in NOS-weighted networks, with longer path length and lower efficiency in BD (*d*=.2; *p*_FDR_=.001), while FA-weighted networks showed no group differences. Males showed lower clustering and local efficiency (*d*=-.3; *p_FDR_*=.0002) than females in NOS-weighted networks, independent of diagnosis, with no sex-by-diagnosis interaction. Age was significantly associated with lower density and efficiency (unweighted), and with path length, local efficiency, and betweenness centrality (NOS-weighted) (*p_FDR_*=[.007-.03]), but no age-by-diagnosis interaction was detected (*p_FDR_*=[.8-.9]). Sorted bar plot of Cohen’s *d* effect sizes for whole-brain network metrics in Figure S2.

**Figure 2.**
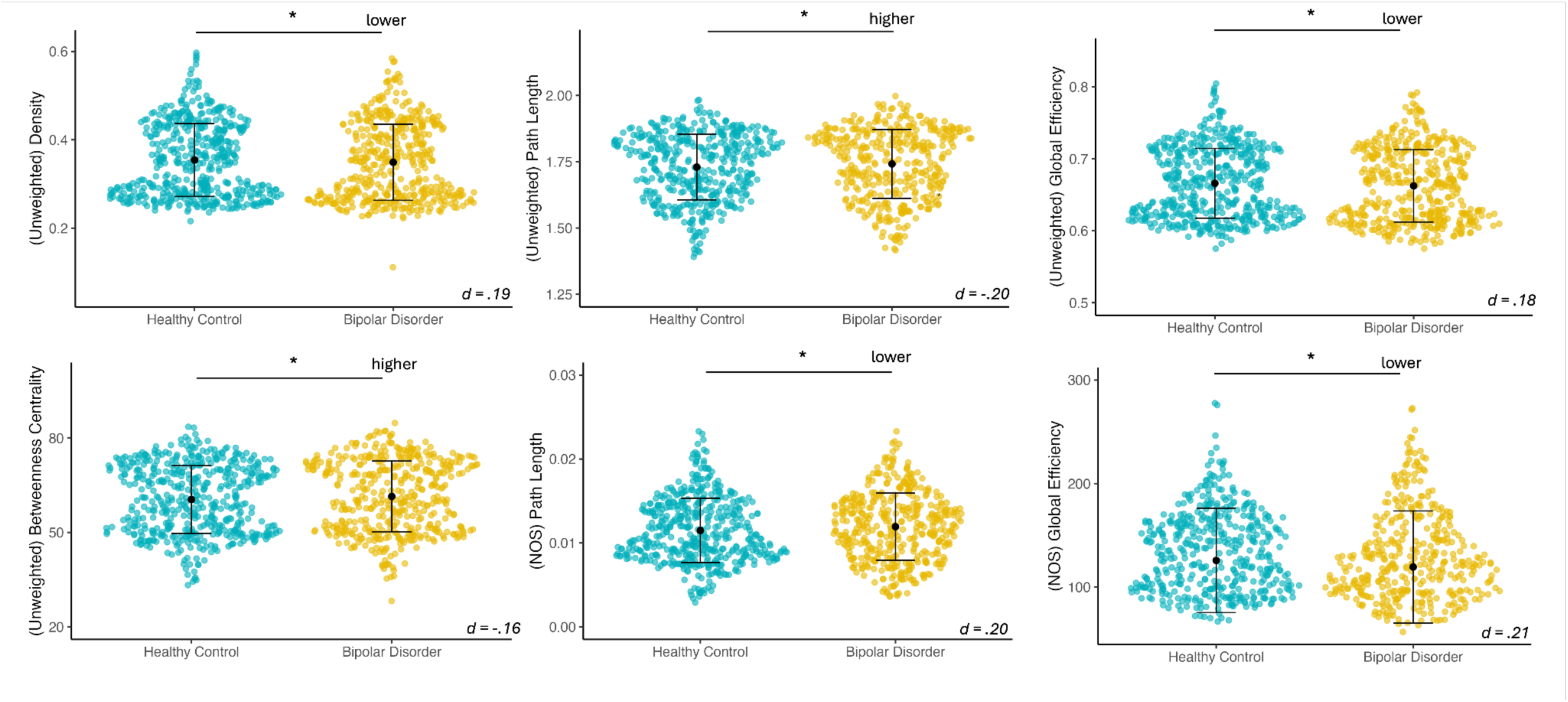
***Whole-brain measures affected in bipolar disorder.*** Global network measures affected in BD, relative to controls. The BD group exhibited dysconnectivity compared to healthy controls across unweighted and NOS-weighted networks. Dysconnectivity was defined by lower global density, longer path length and lower global efficiency; bars represent mean±SD, raw means are plotted. NOS = number of streamlines. \**p_FDR_ <.05*.

**Table 2.**
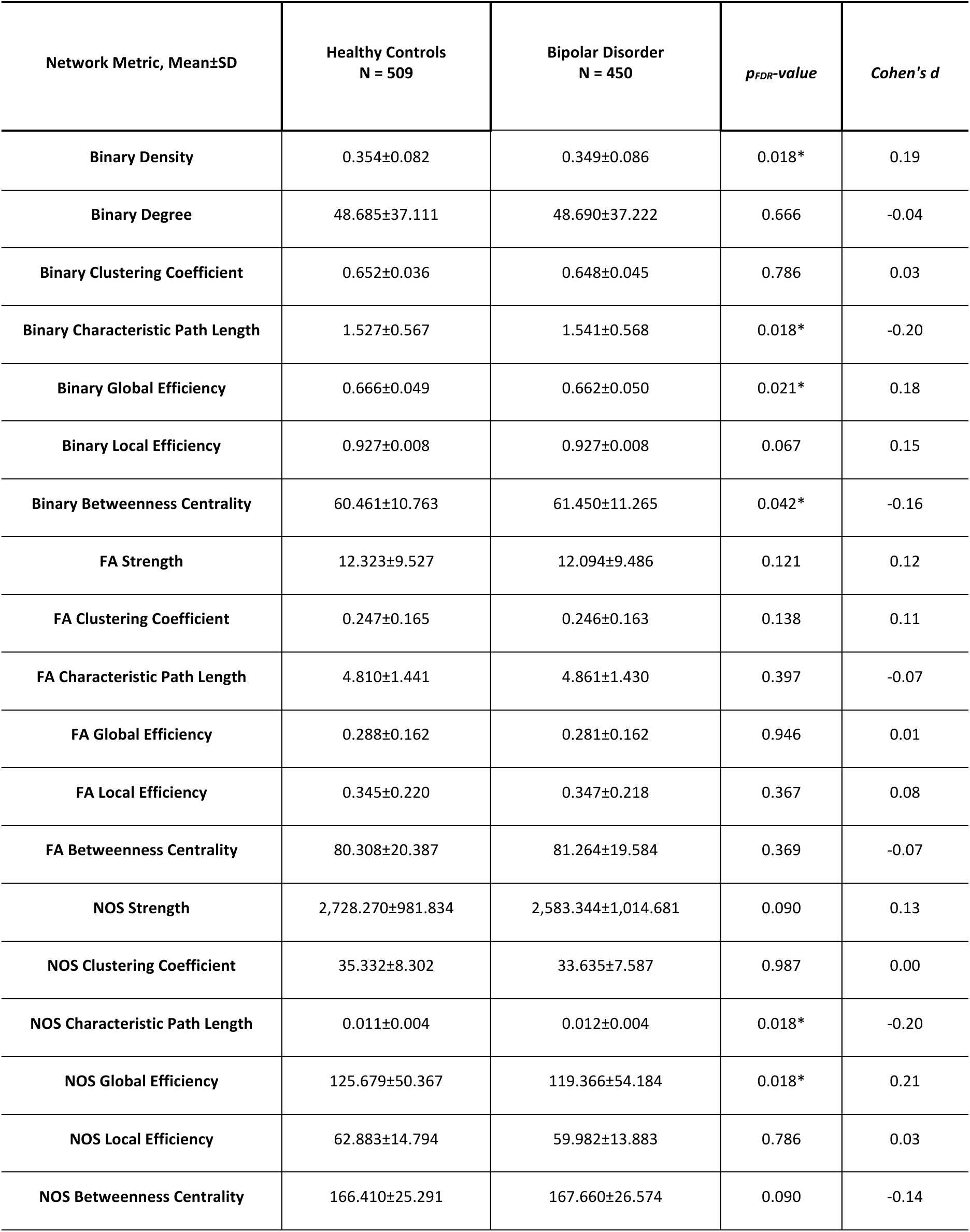
Global Network Measures Across Unweighted and Weighted Networks. Measures are shown across unweighted (binary) networks, with diagnostic group differences in global density, path length, efficiency, and betweenness centrality; weighted networks; at \**p_FDR_* < .05. FA=Fractional Anisotropy. NOS=number of streamlines.

| Network Metric, Mean±SD | Healthy Controls<br>N = 509 | Bipolar Disorder<br>N = 450 | *p<sub>FDR</sub>*-value | Cohen's *d* |
| --- | --- | --- | --- | --- |
| Binary Density | 0.354±0.082 | 0.349±0.086 | 0.018* | 0.19 |
| Binary Degree | 48.685±37.111 | 48.690±37.222 | 0.666 | -0.04 |
| Binary Clustering Coefficient | 0.652±0.036 | 0.648±0.045 | 0.786 | 0.03 |
| Binary Characteristic Path Length | 1.527±0.567 | 1.541±0.568 | 0.018* | -0.20 |
| Binary Global Efficiency | 0.666±0.049 | 0.662±0.050 | 0.021* | 0.18 |
| Binary Local Efficiency | 0.927±0.008 | 0.927±0.008 | 0.067 | 0.15 |
| Binary Betweenness Centrality | 60.461±10.763 | 61.450±11.265 | 0.042* | -0.16 |
| FA Strength | 12.323±9.527 | 12.094±9.486 | 0.121 | 0.12 |
| FA Clustering Coefficient | 0.247±0.165 | 0.246±0.163 | 0.138 | 0.11 |
| FA Characteristic Path Length | 4.810±1.441 | 4.861±1.430 | 0.397 | -0.07 |
| FA Global Efficiency | 0.288±0.162 | 0.281±0.162 | 0.946 | 0.01 |
| FA Local Efficiency | 0.345±0.220 | 0.347±0.218 | 0.367 | 0.08 |
| FA Betweenness Centrality | 80.308±20.387 | 81.264±19.584 | 0.369 | -0.07 |
| NOS Strength | 2,728.270±981.834 | 2,583.344±1,014.681 | 0.090 | 0.13 |
| NOS Clustering Coefficient | 35.332±8.302 | 33.635±7.587 | 0.987 | 0.00 |
| NOS Characteristic Path Length | 0.011±0.004 | 0.012±0.004 | 0.018* | -0.20 |
| NOS Global Efficiency | 125.679±50.367 | 119.366±54.184 | 0.018* | 0.21 |
| NOS Local Efficiency | 62.883±14.794 | 59.982±13.883 | 0.786 | 0.03 |
| NOS Betweenness Centrality | 166.410±25.291 | 167.660±26.574 | 0.090 | -0.14 |

### Permutation-based subnetwork analysis

Edge-level analysis identified a differentially connected subnetwork (FA-weighted) in BD, relative to controls (*t*>1.5, *d=.2; p*_FWE_=.0002) involving 27 structural dysconnections predominantly between and within limbic and basal ganglia nodes, and cerebellar connections via limbic system nodes (Figure 3, Table S9). A differentially connected subnetwork (NOS-weighted) was also seen for BD relative to controls (*t*>10, *d=.5; p*_FWE_=.005) involving 37 structural dysconnections encompassing connections between default-mode/salience network nodes, and fronto-limbic system and basal ganglia nodes (Figure 3, Table S10). No significant (FA/NOS-weighted) weaker/stronger subnetwork was identified when comparing males to females, or when we tested for a diagnosis-by-sex interaction.

**Figure 3.**
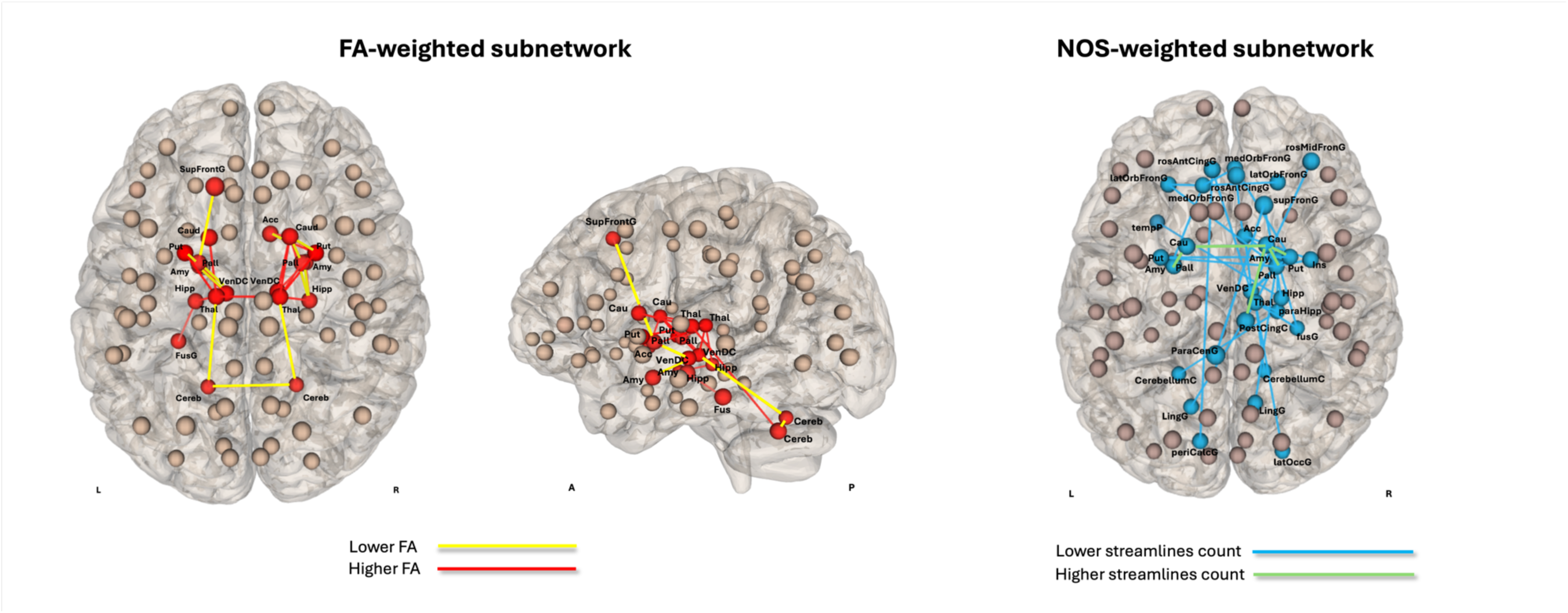
Subnetwork Graph Components Showing Altered FA-weighted and NOS-weighted Connectivity in BD. The left panel displays a network of altered FA-weighted connectivity in BD compared to controls, with the most significant differences observed between and within the basal ganglia and limbic connections (*t* > 1.5, *p*_FWE_ = .0002, *d* = .2). Connections with lower FA-weighted connectivity are shown in yellow, while those with higher FA-weighted connectivity are depicted in red. The right panel illustrates altered NOS-weighted connectivity in BD versus controls (*t* > 10, *p*_FWE_ = .005, *d* = .5), primarily involving the default-mode/salience network regions, fronto-limbic system, and basal ganglia connections. Those connections with lower NOS-weighted connectivity are depicted in blue, while those with higher NOS-connectivity are displayed in green. SupFrontG = Superior Frontal Gyrus; Cau = Caudate; Amy = Amygdala; Hipp = Hippocampus; Thal = Thalamus; FusG = Fusiform Gyrus; Cereb = Cerebellum; VenDC = Ventral diencephalon; Acc = Accumbens; Put = Putamen; Pall = Pallidum; rosAntCingG = Rostral Anterior Cingulate Gyrus; medOrbFronG = Medial Orbitofrontal Gyrus; rosMidFronG = Rostral Middle Frontal Gyrus; supFronG = Superior Frontal Gyrus; tempP = Temporal Pole; Ins = Insula; paraHipp = Parahippocampal Gyrus; PostCingG = Posterior Cingulate Gyrus; LingG = Lingual Gyrus; periCalcG = pericalcarine Cortex; latOccG = Lateral Occipital Gyrus.

### Clinical associations within the Bipolar Disorder Group

At the whole-brain level, longer illness duration (mean±SD: 16±11 years) was associated with lower network density, efficiency, and longer path length (Figure S3a). Later illness onset (mean±SD: 21±9 years) was linked to longer path length and poorer efficiency (Figure S3b). A history of psychosis (N=116) was associated with lower density, longer path length, and higher betweenness centrality compared to those without a psychosis diagnosis (N=112) (Figure S3c). All associations remained significant after adjusting for medication use, apart from illness duration effects on centrality. The number of manic or depressive episodes showed no significant associations with global metrics (*p_FDR_*>.05). Statistical details on clinical associations are provided in Table S11.

In subnetwork analyses, longer illness duration was linked to lower FA-weighted connectivity between the right hippocampus and amygdala, and lower NOS-weighted connectivity between the right cerebellum and thalamus (Figure S3d; Table S11). Later illness onset was associated with lower NOS in connections involving the right cerebellum–thalamus, right amygdala–accumbens, right pallidum–insula, amygdala–fusiform, and left amygdala–medial orbitofrontal cortex (Figure S3d). Greater number of manic episodes was associated with higher NOS-weighted connectivity involving the right anterior middle frontal cortex-ventral diencephalon area, while depressive episodes and psychosis history showed no significant subnetwork associations (*p_FDR_*>.05). Results remained significant after adjusting for medication use. No connectivity (whole-brain and subnetwork) differences were observed between diagnostic BD subtypes I (N=201) and II (N=42).

### Treatment Associations

In those with BD, antidepressant use at time of scan was associated with lower network density, longer path length, lower efficiency, and higher betweenness centrality at the whole-brain level, even after adjusting for illness severity and concurrent medications (Figure S4). No significant associations were observed for lithium, anticonvulsants, or antipsychotics on global network measures (*p_FDR_*>.05). At the subnetwork level, anticonvulsant use was associated with higher NOS-weighted connectivity between the right pallidum and both the right ventral diencephalon and superior frontal cortex, and lower connectivity between the right putamen and medial orbitofrontal cortex (Figure S4). Additional patterns of lower NOS-weighted connectivity were observed between the right thalamus and the fusiform, lingual, and lateral occipital cortices (Figure S4). However, after accounting for illness severity, the associations previously observed with anticonvulsant use were no longer significant. No significant FA/NOS-weighted subnetwork associations were found for lithium, antidepressants, or antipsychotics.

Use of serotonin (5HT) reuptake inhibitors (NbN4) was associated with lower density, longer path length, lower efficiency, and higher betweenness centrality at the whole-brain level, even after adjusting for illness course severity measures and other NbN-based treatment categories (Figure S4). 5HT-reuptake inhibitor use was also linked to lower FA-weighted connectivity between the right thalamus and hippocampus (Figure S4), independent of depressive episodes and other medication classes, but not manic episodes. Use of dopamine, serotonin, and noradrenaline receptor antagonists (NbN2) was associated with lower NOS-weighted connectivity between the left and right middle orbitofrontal gyri, even after accounting for illness course severity scores and other NbN categories (Figure S4). Statistical details on traditional and NbN-classified medication effects are provided in Table S11.

## Discussion

Our study represents the largest analysis of brain network organization in BD to date, analyzing 450 predominantly euthymic individuals with BD relative to 509 controls across 16 international cohorts. BD was associated with altered whole-brain connectivity, and regional dysconnectivity within fronto-limbic and basal ganglia pathways and networks linking default-mode, salience, and basal ganglia regions. Illness severity measures further modulated these alterations: longer illness duration and later onset were associated with more pronounced global and subnetwork disruptions, a history of psychosis with greater deviations in global network features, and a greater number of manic episodes with higher fronto-limbic connectivity. Pharmacological treatments showed distinct, class-specific associations: antidepressant use, particularly selective serotonin reuptake inhibitors (SSRIs), was linked to lower global network integration (lower density and efficiency), longer paths, higher centrality, and reduced microstructural organization between key limbic regions, while anticonvulsants and antipsychotics were associated with alterations in basal ganglia–mediated emotion regulation and cognitive control circuits, including fewer streamlines between frontal cortices.

Our findings reveal a subtle but reliable pattern of whole-brain dysconnectivity in BD across key measures of network integration and segregation (Figure 2), with the largest group differences emerging for characteristic path length and global efficiency. The magnitude and direction of these effects were comparable to those reported in large-scale meta-analytic work (Favre et al., 2019), reinforcing the presence of modest yet consistent disruptions in global network topology. Lower global density and efficiency indicate weaker large-scale integration, while longer paths and elevated betweenness centrality suggest that information flow becomes less direct and more reliant on a limited set of hubs. This configuration reflects a compromised topological organization in BD, one in which fewer hubs carry a disproportionate share of network communication. Clinically, reduced integration and hub overreliance may contribute to cognitive and affective disturbances (Van den Heuvel & Sporns, 2013) and increase vulnerability to destabilization during stress or mood-state transitions, consistent with evidence linking compromised connectivity to episode onset (Sha et al., 2019; Adler et al., 2006; Strakowski et al., 2012). These findings align with prior reports of reduced clustering and efficiency and altered hemispheric or regional connectivity (Leow et al., 2013; Gadelkarim et al., 2014; O’Donoghue et al., 2017; Roberts et al., 2018; Wang et al., 2018), supporting a convergent pattern of global topological abnormalities across studies. We also observed sex-related differences, with males showing lower clustering and local efficiency than females, consistent with prior work on sex-specific connectivity (Allen et al., 2011; Sun et al., 2015) and reports of sex-linked cognitive variation in BD (Suwalska & Łojko, 2014), though these differences did not extend to subnetwork connectivity in our sample.

Whole-brain alterations in BD were driven primarily by dysconnectivity within two subnetworks:

1. fronto-limbic and basal ganglia pathways, including posterior cerebellar projections, and (2) a network linking default-mode, salience, fronto-limbic, and basal ganglia regions (Figure 3). The involvement of default-mode and salience networks in BD points to potential dysfunction in self-referential processing and attentional control (Zovetti et al., 2020), while across both subnetworks the most pronounced effects involved basal ganglia connections, underscoring their central role in BD-related structural disruption and in emotion regulation, reward processing, and cognitive control. Differences between FA-weighted and streamline-weighted networks suggest complementary microstructural (e.g., axonal density, myelination) and macrostructural (tract-level) alterations (Jones et al., 2013; Yeh et al., 2018). The co-occurrence of lower and higher connectivity within these subnetworks is consistent with a pattern of both disruption and compensatory reorganization, in which loss of WM microstructure in core affective pathways may be offset by increased reliance on alternative routes to maintain communication. These divergent patterns indicate that BD involves not only structural disruption but also dynamic, potentially compensatory reorganization aimed at preserving communication across affected circuits. Appreciating these network-level adaptations highlights the need to study both structural and dynamic processes and motivates the search for biomarkers that index these compensatory mechanisms, with the potential to guide future circuit-based interventions in BD. Ongoing efforts within ENIGMA are beginning to bridge such MRI-derived patterns with underlying biological gradients, including cell-type–specific gene expression and neurotransmitter profiles (Patel et al., 2021; Park et al., 2022). The anatomical distribution of effects aligns with prior BD findings, particularly involving the anterior limb of the internal capsule, uncinate fasciculus, ventral amygdalo-striatal projections, cingulum, fornix, and corpus callosum; tracts that link basal ganglia nodes with limbic, frontal, and diencephalic regions implicated in affective regulation (Leow et al., 2013; Ajilore et al., 2015; O’Donoghue et al., 2017; Roberts et al., 2018). Consistent with the largest multi-center ENIGMA-BD DTI study (Favre et al., 2019), which reported widespread FA reductions with strongest effects in the corpus callosum and cingulum, our multisite tractography and network-level approach shows how these regional abnormalities converge into systems-level alterations. This shift from isolated regions to distributed networks is crucial, as converging evidence suggests that psychiatric disorders arise from disturbances in network organization (Sporns, 2011; Bassett & Sporns, 2017), with direct implications for developing circuit-level, mechanism-based interventions.

Our findings also highlight the salience network’s role in BD, particularly its integration with subcortical regions involved in interoception and visceromotor control (Uddin et al., 2011). Within this network, the anterior cingulate cortex (ACC) occupies a strategic connector-hub position linking the amygdala, orbitofrontal cortex, olfactory cortex, and temporal regions critical for emotion regulation and homeostasis (Mesulam & Mufson, 1982). Dynamic instabilities in interoceptive networks may underlie the maladaptive responses to emotional stimuli observed in BD (Perry et al. 2018), and consequent aberrant perception of emotional stimuli as increasingly salient (Nabulsi et al. 2022). The elevated betweenness centrality observed in BD, together with prior evidence of altered connector-hub organization (Nabulsi et al., 2019; Nabulsi et al., 2022), suggests increased reliance on these hubs to compensate for lower structural connectivity within core affective and interoceptive pathways. Such hub overreliance may reflect compensatory engagement of connector hubs that support the integration of external stimuli with self-referential processing–a process known to be disrupted in BD and other mood disorders (Barch, 2005; Phillips et al., 2008; Menon, 2015; Perry et al., 2018)–and may be key to the regulation of emotional experiences in BD.

Clinical features were significantly associated with the degree of topological arrangement in BD (Figure S3). Older individuals with BD showed greater alterations in amygdala–hippocampal connectivity, consistent with cumulative disease burden, and longer illness duration was linked to widespread reductions in network integration across large-scale systems. Although our cross-sectional design limits causal inference, these patterns align with evidence that chronic neuroinflammatory processes and glial activation contribute to progressive WM abnormalities in BD (Benedetti et al., 2016). Later illness onset was associated with more pronounced global topological alterations, particularly within cerebello-thalamic and fronto-limbic–basal ganglia pathways—including reduced connectivity along uncinate fasciculus–related regions—suggesting increased vulnerability in individuals who do not receive early intervention (Weathers et al., 2018). A history of psychosis was linked to a more fragmented and inefficient global network, consistent with greater reliance on a limited set of high-betweenness nodes to compensate for broader connectivity deficits. In contrast, a greater number of manic episodes was associated with higher connectivity between the anterior middle frontal cortex and subcortical regions inferior to the thalamus (hypothalamus, mammillary bodies, subthalamic nuclei, substantia nigra), in line with evidence that mania-induced neural plasticity may alter fronto-limbic circuitry via dopaminergic and glutamatergic mechanisms (Manji et al., 2003; Abé et al., 2021; Strakowski et al., 2012; Houenou et al., 2007). The absence of similar effects for depressive episodes suggests distinct mood-state–dependent mechanisms, although our predominantly euthymic sample and cross-sectional design may limit our ability to detect acute or residual mood-related structural changes (Vieta & De Prisco, 2024). These results demonstrate that illness-related factors—including duration, onset age, psychosis history, and episode frequency—must be modeled to understand BD network pathology, and they motivate longitudinal, deeply phenotyped studies to determine whether observed alterations reflect illness burden, compensatory reorganization, or cumulative manic effects, a distinction with direct relevance for early, mechanism-guided intervention strategies.

Antidepressant use, particularly selective serotonin reuptake inhibitors (SSRIs), was associated with lower connectivity within limbic circuits involving the thalamus and hippocampus and with reduced global network integration. This pattern raises the possibility that, while SSRIs may help stabilize mood, their mechanism of action could differentially affect WM connectivity in regions already vulnerable to illness severity in BD. This interpretation should be made cautiously: medication use was not randomly assigned, and although our models accounted for symptom severity and number of depressive episodes, residual confounding and unmeasured aspects of illness chronicity may persist, particularly among individuals more likely to be prescribed SSRIs across ENIGMA-BD samples. ENIGMA studies of major depressive disorder found cortical and hippocampal differences in medicated patients but no WM microstructural changes (Schmaal et al., 2020), suggesting that antidepressant-related WM alterations may be specific to BD and influenced by illness burden. Anticonvulsant use produced mixed connectivity changes within emotion-regulation and cognitive-control circuits, overlapping with BD social-cognition networks (Sha et al., 2019) and regions sensitive to anticonvulsant-related cognitive effects (Meador, 2003). These associations disappeared after adjusting for illness severity, highlighting the challenge of separating medication effects from disease burden and the need for prospective, dose-resolved longitudinal designs. Antipsychotic use was linked to lower connectivity between the medial orbitofrontal cortex and other frontal regions, a pattern that may reflect either therapeutic modulation of limbic–frontal integration or potential cognitive side effects; distinguishing these possibilities will require longitudinal and dose-resolved data. Lithium and other mood stabilizers did not show clear associations with network connectivity in our sample. This absence may reflect lithium’s distinct intracellular signaling mechanisms—which differ from the receptor- or ion-channel–based actions of SSRIs, antipsychotics, and anticonvulsants—and is consistent with its proposed neuroprotective influence (Hibar et al., 2016; 2017), potentially indicating a stabilizing effect on WM architecture even if not significant here. The relative preservation of network organization in lithium-treated individuals offers a translational distinction between medications that maintain structural connectivity and those associated with reduced integration, suggesting that connectomic markers could help differentiate treatments by their network impact. Whether this reflects methodological limitations or a true mechanistic distinction from other psychotropics requires further study. Overall, although longitudinal studies remain necessary to disentangle medication effects from illness progression, our findings support the broader notion that psychotropic medications influence network-level organization in circuits aligned with their mechanistic targets, raising important questions about whether these changes are adaptive, maladaptive, or necessary trade-offs in clinical management.

Our findings demonstrate the feasibility and scientific value of large-scale, multisite connectome analyses in BD using harmonized diffusion MRI and standardized anatomical pipelines. In the largest BD connectome study to date, we identified subtle but robust alterations in emotion-regulation and reward-related networks that were further shaped by illness burden and pharmacological treatment. We report differential effects based on the mechanism of action of medications, with antidepressant-related alterations in limbic connectivity illustrating the importance of accounting for treatment exposure and of developing more sophisticated approaches for modeling medication effects when interpreting network abnormalities. More broadly, these results illustrate how network-based approaches capture system-level disruptions not apparent in regional analyses and provide a framework for disentangling pharmacological influences from core neurobiological features of BD. Continued collaborative efforts at this scale will be critical for biomarker development and for advancing more precise, circuit-informed treatment strategies in psychiatric disorders.

## Acknowledgments

D.M.C., L.N., G.M.P., C.M.D., B.H.: The participants and the support of the Wellcome-Trust Health Research Board Clinical Research Facility and the Centre for Advanced Medical Imaging, St. James Hospital, Dublin, Ireland; DS, HST, FS, FH: SAMRC, University Research Committee, University of Cape Town and South African funding bodies National Research Foundation (SANRF) and Medical Research Council (SAMRC); Marinka MG Koenis for data support (Yale/Olin Cohort).

## Contribution

**Cohort PI:** O.A.A., M.Bel., M.Ber., P.B., D.M.C., C.R.K.C., U.D., D.C.G., D.G., B.Haa.,T.H., J.H., F.M.H., T.K., C.M., P.B.M., P.M., M.L.P., M.P., E.P.-C., E.V., M.W., H.T. **Project leader:** L.N., D.M.C. **Core analysis/writing group:** D.M.C., C.R.K.C., M.J.Y.K., L.N., P.M.T. **Clinical data collection** M.Bel., M.Ber., T.B., E.J.C.-R., O.D., U.D., P.F., K.F., B.Hal., J.H., F.M.H., A.J., A.K., E.J.L., F.M.M., C.M., G.M., S.M., P.B.M., L.N., S.O., M.L.P., M.P., A.P., J.Ra., J.Re., G.R., A.L.R., R.S., F.Sc., F.St., O.S., C.S., H.S.T., L.T., F.T.-O., P.L.U., G.M., D.C.G., H.T. **Imaging data collection** M.Bel., M.Ber., T.B., E.J.C.-R., O.D., U.D., P.F., K.F., J.H., F.M.H., A.J., A.K., F.M.M., G.M., C.M., S.M., P.B.M., L.N., S.O., M.L.P., M.P., A.P., J.Ra., J.Re., G.R., A.L.R., R.S., F.Sc., F.St., O.S., C.S., H.S.T., L.T., F.T.-O., D.C.G., J.Rep., B.S. **Manuscript revision/editing** N.A., O.A.A., M.Bel., C.R.K.C., O.D., U.D., P.F., D.C.G., T.H., J.H., F.M.H., N.J., M.J.Y.K., C.M., P.B.M., L.N., M.P., J.Ra., G.R., D.J.S., F.St., B.S., S.I.T., P.M.T., N.P., H.T., J. Rep.

