## Supplemental Material for "Structural Brain Network Alterations in Relation to Treatment and Illness Severity in Bipolar Disorder"

### Supplementary Material

#### *Image acquisition and processing*

Contributing sites shared their raw T1-weighted and diffusion-weighted (dMRI) scans with the main project team. Details on acquisition parameters for each site are detailed in Table S3. Structural MR images were visually inspected before and after processing to ensure the accuracy of cortico-subcortical parcellation and segmentation, particularly examining gray and white matter boundaries. A probabilistic approach was employed to map subject-specific cortico-subcortical brain networks encompassing 34 cortical and 9 subcortical brain regions on both sides and including the cerebellum (FreeSurfer v5.3.0) (Fischl 2012). This mapping was based on the Desikan-Killiany atlas (Desikan et al. 2006) applied to any T1-weighted image, covering a total of 86 regions. Parcellation/segmentation from T1-weighted images was evaluated using standard ENIGMA protocols (https://enigma.ini.usc.edu/protocols/imaging-protocols/;https://github.com/ENIGMA-git/ENIGMA-FreeSurfer-protocol). Diffusion MR images were corrected for subject motion, including rotating the b-matrix and addressing eddy-current distortions (ExploreDTI v4.8.6) (Leemans and Jones 2009). Images were carefully inspected for artifacts, head motion, signal dropout, eddy-current-induced distortion and partial volume effects using in-house scripts. To resolve crossing fibers within voxels we employed a deterministic constrained spherical deconvolution (CSD) algorithm (*lmax* = 6) (ExploreDTI v4.8.6) (Tournier et al. 2007; Jeurissen et al. 2014). Diffusion eigenvector estimation was performed with the robust estimation of tensor by outlier rejection (RESTORE) method (Chang et al. 2005). Fiber tracking initiated in each voxel, followed a 1 mm step size and a 2 $mm^3$ seed point resolution, maintained a curvature threshold greater than 30°, covered lengths between 20 and 300 mm, and terminated at a minimum fractional anisotropy (FA) of 0.2. For each subject, whole-brain tractography maps were subsequently used alongside the parcellated T1 labels to generate individual (86 × 86) undirected connectivity matrices (Figure 1) (ExploreDTI v4.8.6). Connectivity matrices were weighted by fractional anisotropy (FA), representing the average FA between two nodes in the network, and by the number of streamlines (NOS), indicating the number of reconstructed trajectories between two nodes. FA and NOS weights are generated and dependent by the tractography step, providing crucial information on the microstructural organization and density of the connections, respectively. Unweighted, or binary, matrices were additionally derived. Topological organization of the networks, both in the weighted and unweighted matrices, was analyzed for both bipolar disorder (BD) and controls.

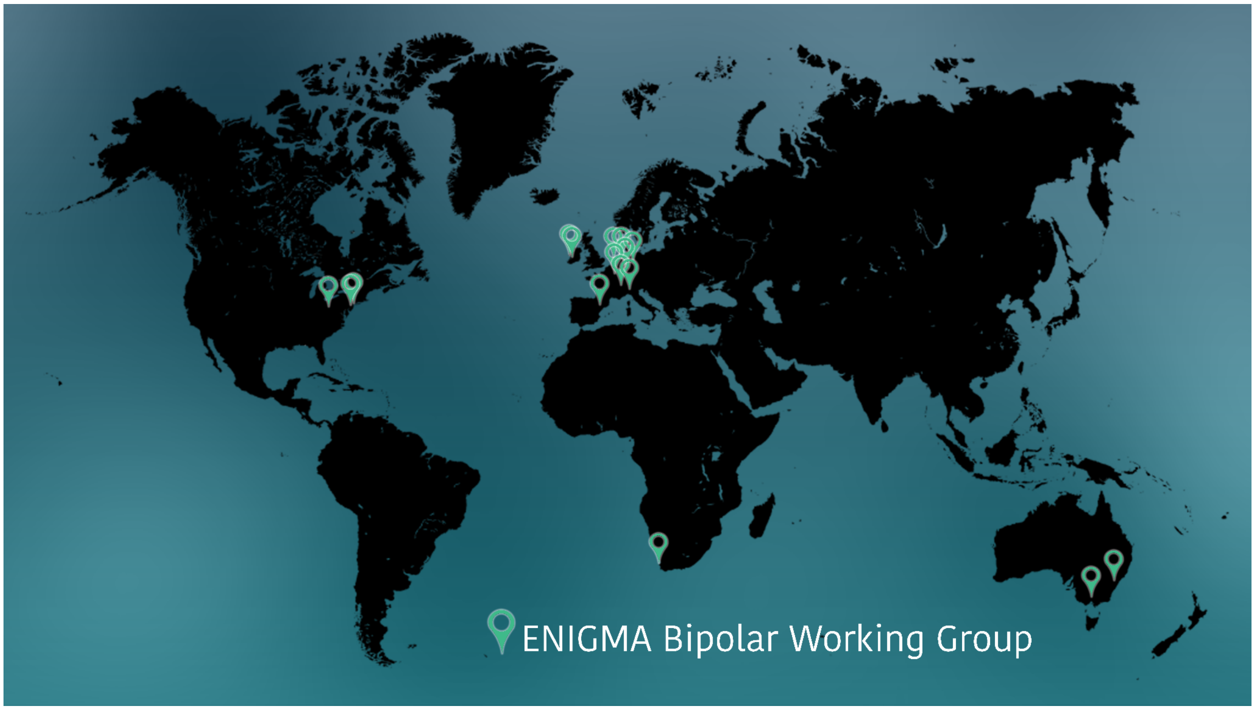


**Figure S1. Global Map of Participating Study Sites.** Sites contributing imaging and clinical data to the study. Participating institutions span multiple countries and reflect the international scope of the collaborative effort.

| | |
|---|---|
| **Lithium,** n / N (%) | |
| On | 134 / 352 (38%) |
| Off | 218 / 352 (62%) |
| NR | 98 |
| **Antipsychotic,** n / N (%) | |
| On | 162 / 359 (45%) |
| Off | 197 / 359 (55%) |
| NR | 91 |
| **Anticonvulsants,** n / N (%**)** | |
| On | 177 / 339 (52%) |
| Off | 162 / 339 (48%) |
| NR | 111 |
| **Antidepressants,** n / N (%) | |
| On | 146 / 353 (41%) |
| Off | 207 / 353 (59%) |
| NR | 97 |
| **NbN1 (Primarily Dopamine Receptor Antagonist),** n / N (%) | |
| On | 18 / 359 (5%) |
| Off | 341 / 359 (95%) |
| NR | 91 |
| **NbN2 (Dopamine and Other (Ser-nor) Monoamine Receptor Antagonists),** n / N (%) | |
| On | 115 / 359 (32%) |
| Off | 244 / 359 (68%) |
| NR | 91 |
| **NbN3 (Dopamine, Serotonin Receptor Partial Agonist/Antagonists),** n / N (%) | |
| On | 21 / 359 (6%) |
| Off | 338 / 359 (94%) |
| NR | 91 |
| **NbN4 (Targeting Serotonin (Reuptake Inhibitors Multimodal),** n / N (%) | |
| On | 63 / 353 (18%) |
| Off | 290 / 353 (82%) |
| NR | 97 |
| **NbN5 (Targeting Serotonin and Other Monoamines with different MOAs),** n / N (%) | |
| On | 34 / 353 (10%) |
| Off | 319 / 353 (90%) |
| NR | 97 |
| **NbN6 (Glutamate, Sodium, Calcium Channel Blockers),** n / N (%) | |
| On | 83 / 339 (24%) |
| Off | 256 / 339 (76%) |
| NR | 111 |
| **NbN7 (Lithium), n / N (%)** | |
| On | 114 / 450 (25%) |
| Off | 336 / 450 (75%) |
| NR | 0 |
| **NbN8 (Valproate), n / N (%)** | |
| On | 52 / 450 (12%) |
| Off | 398 / 450 (88%) |
| NR | 0 |
| **NbN9 (Others), n / N (%)** | |
| On | 59 / 450 (13%) |

| | |
|---|---|
| Off | 391 / 450 (87%) |
| NR | 0 |

**Table S1. Traditional-indication based nomenclature and Neuroscience-based Nomenclature (NbN) categories samples.** This table presents the demographics of participants categorized according to the traditional medication naming convention the Neuroscience-based Nomenclature (NbN) framework. n/N = Sample/Total Sample Size. MOA = Mechanism Of Action. NR= not reported.

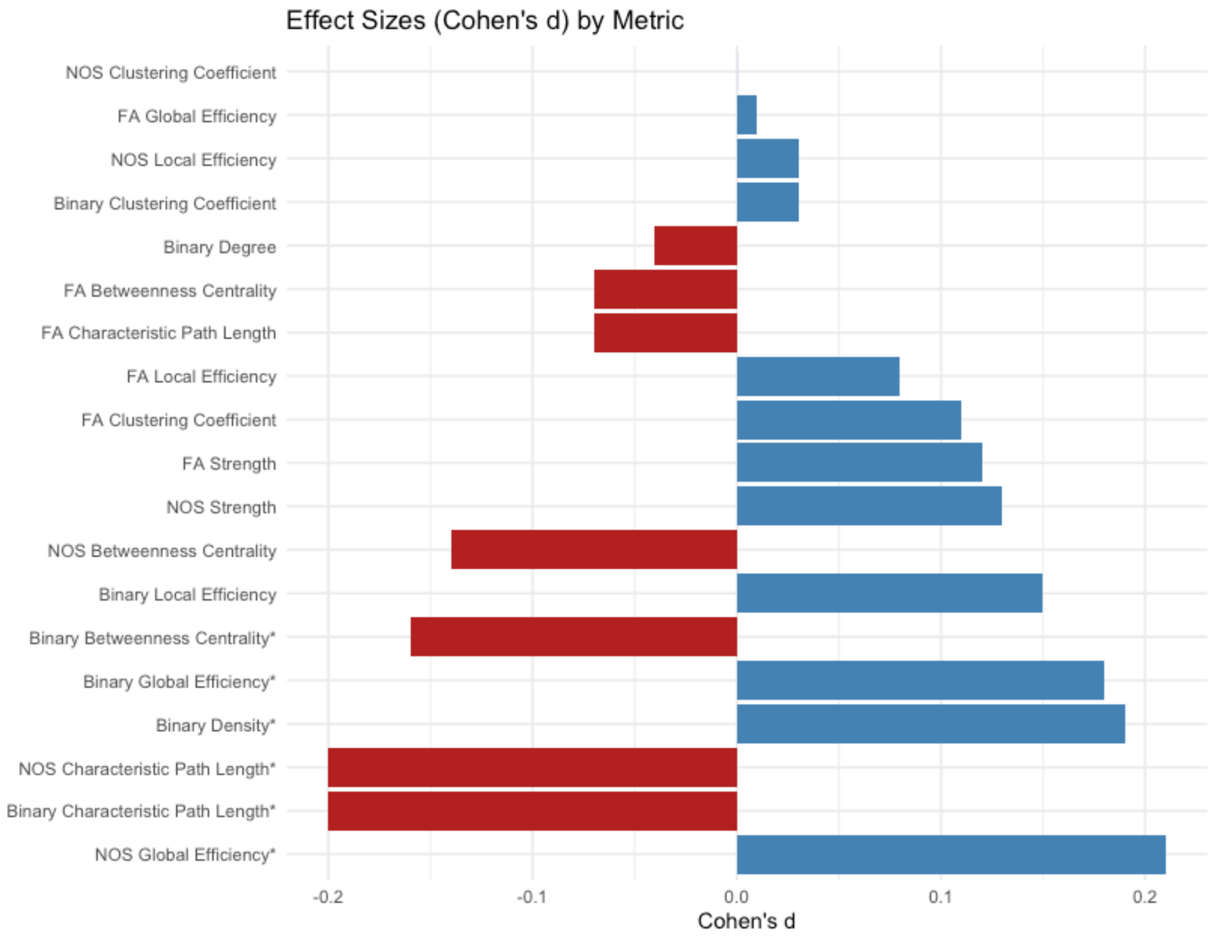


**Figure S2. Sorted bar plot of Cohen's *d* effect sizes for whole-brain network metrics in bipolar disorder *versus* healthy controls.** Effect sizes represent the standardized mean difference (Cohen's *d*) between diagnostic groups for each network measure. Metrics are sorted by the absolute magnitude of effect size to highlight the strongest group differences. *$p_{FDR}$<.05

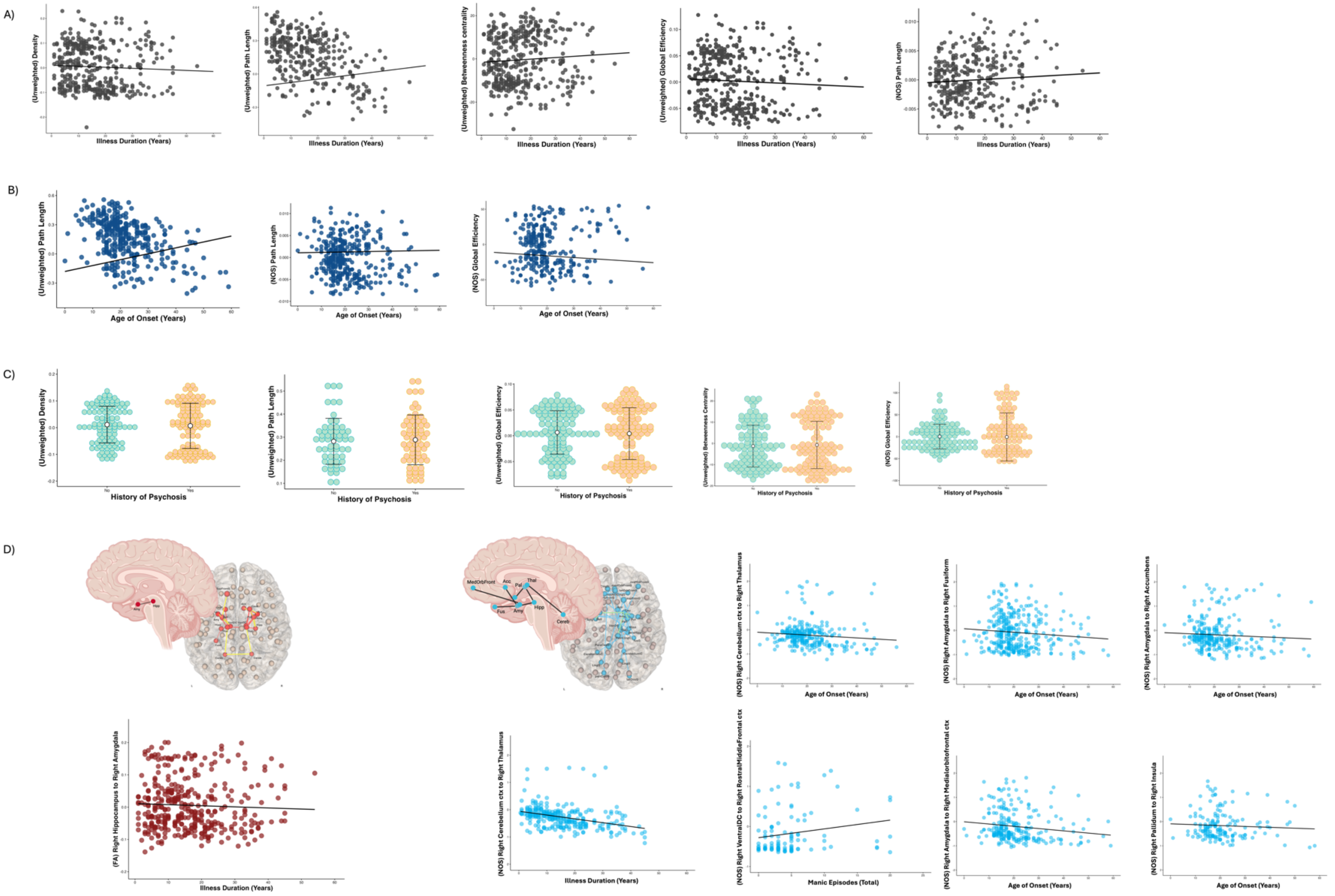


**Figure S3. Whole-Brain and Subnetwork Changes Associated with Clinical Variables in Bipolar Disorder.** Whole-brain network changes in the bipolar disorder (BD) group are associated with (A) illness duration, which correlates with lower network density, poorer efficiency, longer path length, and higher betweenness centrality; (B) age of onset, linked to poorer network features such as longer path length and reduced global efficiency; and (C) history of psychosis, associated with decreased connectivity density, global efficiency, longer path length, and increased betweenness centrality. Additionally, (D) subnetwork changes indicate that longer illness duration is associated with lower FA-weighted connectivity between the right hippocampus and right amygdala, and lower NOS-weighted connectivity between the right cerebellum and right thalamus. A later onset of illness associated with lower NOS-weighted connectivity among several connections, including between the right cerebellum and right thalamus, right amygdala and right accumbens area, right pallidum and right insula, right amygdala and right fusiform, as well as between the left amygdala and left medial orbitofrontal cortex. The number of manic episodes was associated with higher NOS-weighted connectivity between the right ventral diencephalon and the right rostralmiddlefrontal cortex.

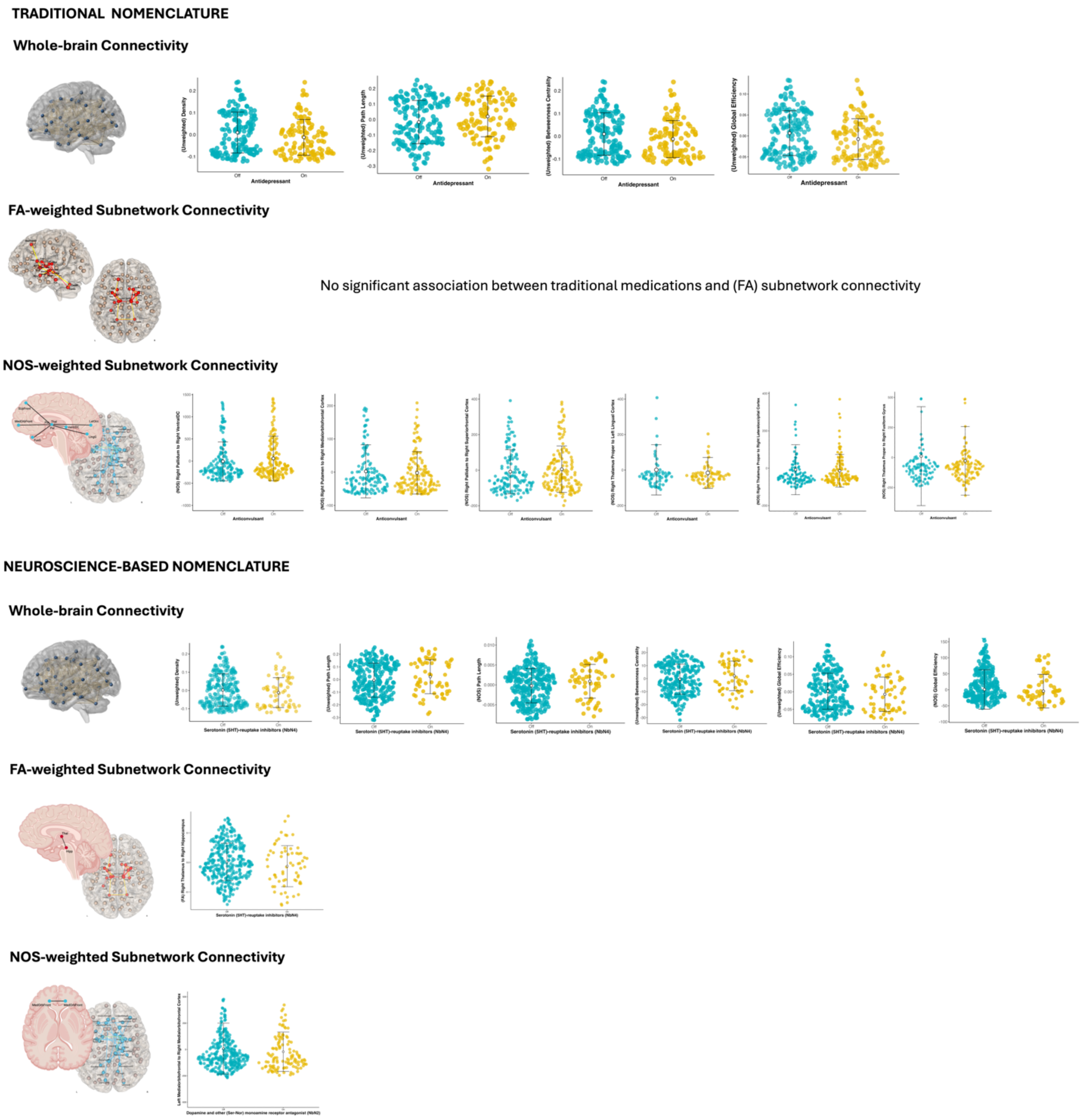


**Figure S4. Associations of Antidepressant, Anticonvulsant and Antipsychotic Use on Whole-Brain and Subnetwork Connectivity in Bipolar Disorder.** Antidepressant use in the BD group was associated with significant changes across whole-brain measures, including lower network density, longer path length, reduced efficiency, and higher betweenness centrality. Lithium, anticonvulsants, and antipsychotics did not show significant associations with whole-brain network metrics. In subnetwork analyses, anticonvulsant use was linked to higher NOS-weighted connectivity between the right pallidum and both the right ventral diencephalon and right superior frontal cortex, while lower NOS-weighted connectivity was observed between the right putamen and the right medial orbitofrontal cortex. Additionally, anticonvulsant use was associated with lower NOS-weighted connectivity between the right thalamus and various occipital and fusiform regions. Using the Neuroscience-based Nomenclature (NbN) framework, 5HT-reuptake inhibitors (NbN4, antidepressants) were associated with similar whole-brain changes, including lower network density, longer path length, reduced efficiency, and higher betweenness centrality. Subnetwork analysis showed lower FA-weighted connectivity between the right thalamus and right hippocampus. Dopamine, serotonin, and noradrenaline receptor antagonists (NbN2, antipsychotics) were associated with lower NOS-weighted connectivity between the left and right middle orbitofrontal gyri. These findings held even after adjusting for symptom severity.

**Table S2. Demographic and Clinical Characteristics of the Study Cohorts.** This table provides detailed demographic and clinical information for the 16 cohorts included in the study. Key demographic variables include age, sex, BD subtype and mood state. Additionally, clinical characteristics including illness duration, age of onset, history of psychosis, number of manic and depressive episodes, and medication use (e.g., antidepressants, antipsychotics, lithium, and anticonvulsants) are shown, offering a comprehensive overview of the clinical profiles relevant to the study. n/N = sample/total sample size; % = percentage.

| Diagnosis, n/N (%) | Barcelona N = 49 | Moodinflame N = 44 | FOR-2017 Munster N = 87 | Pittsburgh N = 62 | Sydney N = 112 | Milano N = 42 | Yale/Odin_33 gradients N = 15 | Yale/Odin_55 gradients N = 127 | Cape Town N = 19 | NUIG-CHRM2 N = 59 | Creteil N = 57 | Deakin N = 68 | NUIG-GBS N = 65 | Grenoble N = 13 | Mannheim N = 51 | FOR-2017 Marburg N = 89 |
|---|---|---|---|---|---|---|---|---|---|---|---|---|---|---|---|---|
| **Healthy Controls** | 17 / 49 (35%) | 23 / 44 (52%) | 71 / 87 (82%) | 21 / 62 (34%) | 60 / 112 (54%) | 30 / 42 (71%) | 11 / 15 (73%) | 52 / 127 (41%) | 10 / 19 (53%) | 38 / 59 (64%) | 36 / 57 (63%) | 28 / 68 (41%) | 32 / 65 (49%) | - | 24 / 51 (47%) | 56 / 89 (63%) |
| **Bipolar Disorder** | 32 / 49 (65%) | 21 / 44 (48%) | 16 / 87 (18%) | 41 / 62 (66%) | 52 / 112 (46%) | 12 / 42 (29%) | 4 / 15 (27%) | 75 / 127 (59%) | 9 / 19 (47%) | 21 / 59 (36%) | 21 / 57 (37%) | 40 / 68 (59%) | 33 / 65 (51%) | 13 / 13 (100%) | 27 / 51 (53%) | 33 / 89 (37%) |
| **Age** | | | | | | | | | | | | | | | | |
| **Mean (SD)** | 39 (9) | 40 (14) | 30 (12) | 33 (8) | 24 (4) | 39 (15) | 33 (11) | 33 (12) | 26 (4) | 41 (13) | 35 (12) | 21 (2) | 40 (10) | 46 (9) | 43 (11) | 44 (12) |
| **Sex** | | | | | | | | | | | | | | | | |
| **Male** | 20 / 49 (41%) | 14 / 44 (32%) | 33 / 87 (38%) | 17 / 62 (27%) | 68 / 112 (61%) | 22 / 42 (52%) | 11 / 15 (73%) | 89 / 127 (70%) | 11 / 19 (58%) | 18 / 59 (31%) | 29 / 57 (51%) | 43 / 68 (63%) | 33 / 65 (51%) | 8 / 13 (62%) | 23 / 51 (45%) | 30 / 89 (34%) |
| **Female** | 29 / 49 (59%) | 30 / 44 (68%) | 54 / 87 (62%) | 45 / 62 (73%) | 44 / 112 (39%) | 20 / 42 (48%) | 4 / 15 (27%) | 38 / 127 (30%) | 8 / 19 (42%) | 41 / 59 (69%) | 28 / 57 (49%) | 25 / 68 (37%) | 32 / 65 (49%) | 5 / 13 (38%) | 28 / 51 (55%) | 59 / 89 (66%) |
| **BD Type, n/N (%)** | | | | | | | | | | | | | | | | |
| **BD 1** | 32 / 32 (100%) | 21 / 21 (100%) | - | 41 / 41 (100%) | 28 / 52 (54%) | NA | NA | NA | 9/9 | 18 / 21 (86%) | 21 | NA | 32 / 32 (100%) | 5 / 11 (45%) | NA | 24 / 32 (75%) |
| **BD 2** | - | - | 1 / 1 (100%) | - | 24 / 52 (46%) | NA | NA | NA | NA | 3 / 21 (14%) | 0 | NA | - | 6 / 11 (55%) | NA | 8 / 32 (25%) |
| **NA, N** | 17 | 23 | 86 | 21 | 60 | 42 | 15 | 127 | 0 | 38 | 57 | 68 | 33 | 2 | 51 | 57 |
| **Mood Phase, n/ N (%)** | | | | | | | | | | | | | | | | |
| **Depressed** | - | - | 5 / 12 (42%) | 12 / 41 (29%) | 2 / 52 (3.8%) | NA | - | 5 / 75 (6.7%) | - | - | 7 / 19 (37%) | NA | 1 / 27 (3.7%) | - | NA | 4 / 33 (12%) |
| **Euthymic** | 32 (100%) | 21 (100%) | 2 / 12 (17%) | 29 / 41 (71%) | 49 / 52 (94%) | NA | 4 / 4 (100%) | 63 / 75 (84%) | 9 / 9 (100%) | 21 / 21 (100%) | 12 / 19 (63%) | NA | 26 / 27 (96%) | 13 / 13 (100%) | NA | 27 / 33 (82%) |
| **Hypomanic** | - | - | 4 / 12 (33%) | - | - | NA | - | - | - | - | - | NA | - | - | NA | - |
| **Manic** | - | - | - | - | - | NA | - | 7 / 75 (9.3%) | - | - | - | NA | - | - | NA | 2 / 33 (6.1%) |
| **Mixed** | - | - | 1 / 12 (8.3%) | - | 1 / 52 (1.9%) | NA | - | - | - | - | - | NA | - | - | NA | - |
| **NA, N** | 17 | 23 | 75 | 21 | 60 | 42 | - | 52 | 10 | 38 | 38 | 68 | 38 | 0 | 51 | 56 |
| **Duration of Illness (Years)** | | | | | | | | | | | | | | | | |
| **Mean (SD)** | 24 (8) | 24 (12) | 14 (10) | 15 (7) | 10 (4) | 18 (14) | NA | 15 (12) | 6 (5) | 18 (10) | 13 (9) | NA | 10 (9) | 17 (8) | 16 (11) | 21 (12) |
| **NA, N** | 17 | 23 | 71 | 21 | 61 | 30 | 15 | 57 | 10 | 41 | 38 | 68 | 35 | 3 | 26 | 56 |
| **Age of Onset (Years)** | | | | | | | | | | | | | | | | |
| **Mean (SD)** | 16 (9) | 21 (8) | 24 (12) | 18 (7) | 15 (3) | 35 (14) | NA (NA) | 19 (8) | 22 (3) | 24 (9) | 23 (7) | NA (NA) | 27 (7) | 30 (9) | 24 (10) | 23 (11) |
| **NA, N** | 17 | 23 | 71 | 21 | 61 | 30 | 15 | 57 | 10 | 41 | 38 | 68 | 35 | 3 | 26 | 56 |
| **Lithium, n/ N (%)** | | | | | | | | | | | | | | | | |
| **Off** | 15 / 32 (47%) | 9 / 21 (43%) | 81 / 87 (93%) | 31 / 41 (76%) | 37 / 52 (71%) | NA | 15 / 15 (100%) | 108 / 126 (86%) | 16 / 19 (84%) | NA | 9 / 20 (45%) | NA | 36 / 55 (65%) | NA | 10 / 16 (63%) | 25 / 34 (74%) |
| **On** | 17 / 32 (53%) | 12 / 21 (57%) | 6 / 87 (6.9%) | 10 / 41 (24%) | 15 / 52 (29%) | 3 / 3 (100%) | NA | 18 / 126 (14%) | 3 / 19 (16%) | 5 / 5 (100%) | 11 / 20 (55%) | NA | 19 / 55 (35%) | NA | 6 / 16 (38%) | 9 / 34 (26%) |
| **NA** | 17 | 23 | 0 | 21 | 60 | 39 | 0 | 1 | 0 | 54 | 37 | 68 | 10 | 13 | 35 | 55 |
| **Antipsychotic, n/ N (%)** | | | | | | | | | | | | | | | | |
| **Off** | 11 / 31 (35%) | 15 / 21 (71%) | 78 / 87 (90%) | 18 / 41 (44%) | 36 / 52 (69%) | 0 / 4 (0%) | 14 / 15 (93%) | 98 / 127 (77%) | 12 / 19 (63%) | NA) | 9 / 20 (45%) | NA | 49 / 53 (92%) | NA | 11 / 11 (100%) | 20 / 34 (59%) |
| **On** | 20 / 31 (65%) | 6 / 21 (29%) | 9 / 87 (10%) | 23 / 41 (56%) | 16 / 52 (31%) | 4 / 4 (100%) | 1 / 15 (6.7%) | 29 / 127 (23%) | 7 / 19 (37%) | 18 / 18 (100%) | 11 / 20 (55%) | NA | 4 / 53 (7.5%) | NA | NA | 14 / 34 (41%) |

| | | | | | | | | | | | | | | | | |
|---|---|---|---|---|---|---|---|---|---|---|---|---|---|---|---|---|
| **NA** | 18 | 23 | 0 | 21 | 60 | 38 | 0 | 0 | 0 | 41 | 37 | 68 | 12 | 13 | 40 | 55 |
| **Anticonvulsants, n/ N (%)** | | | | | | | | | | | | | | | | |
| **Off** | 15 / 32 (47%) | 10 / 21 (48%) | 85 / 87 (98%) | 33 / 62 (53%) | 19 / 48 (40%) | NA | 13 / 15 (87%) | 87 / 126 (69%) | 13 / 19 (68%) | NA) | 10 / 20 (50%) | NA | 48 / 55 (87%) | NA | 6 / 15 (40%) | 66 / 75 (88%) |
| **On** | 17 / 32 (53%) | 11 / 21 (52%) | 2 / 87 (2.3%) | 29 / 62 (47%) | 29 / 48 (60%) | 6 / 6 (100%) | 2 / 15 (13%) | 39 / 126 (31%) | 6 / 19 (32%) | 1 / 1 (100%) | 10 / 20 (50%) | NA | 7 / 55 (13%) | NA | 9 / 15 (60%) | 9 / 75 (12%) |
| **NA** | 17 | 23 | 0 | 0 | 64 | 36 | 0 | 1 | 0 | 58 | 37 | 68 | 10 | 13 | 36 | 14 |
| **Antidepressants, n/ N (%)** | | | | | | | | | | | | | | | | |
| **Off** | 25 / 31 (81%) | 15 / 21 (71%) | 80 / 87 (92%) | 22 / 41 (54%) | 28 / 52 (54%) | NA | 12 / 15 (80%) | 87 / 126 (69%) | 19 / 19 (100%) | NA | 13 / 20 (65%) | NA | 50 / 55 (91%) | NA | 6 / 15 (40%) | 23 / 34 (68%) |
| **On** | 6 / 31 (19%) | 6 / 21 (29%) | 7 / 87 (8.0%) | 19 / 41 (46%) | 24 / 52 (46%) | 6 / 6 (100%) | 3 / 15 (20%) | 39 / 126 (31%) | NA | 5 / 5 (100%) | 7 / 20 (35%) | NA | 5 / 55 (9.1%) | NA | 9 / 15 (60%) | 11 / 34 (32%) |
| **NA** | 18 | 23 | 0 | 21 | 60 | 36 | 0 | 1 | 0 | 54 | 37 | 68 | 10 | 13 | 36 | 55 |
| **History of Psychosis, n/ N (%)** | | | | | | | | | | | | | | | | |
| **No** | 7 / 31 (23%) | NA | NA | NA | 37 / 52 (71%) | NA | NA | 38 / 64 (59%) | 0 / 9 (0%) | 19 / 21 (90%) | NA | NA | 32 / 55 (58%) | 8 / 11 (73%) | NA | NA |
| **Yes** | 24 / 31 (77%) | NA | NA | NA | 15 / 52 (29%) | NA | NA | 26 / 64 (41%) | 9 / 9 (100%) | 2 / 21 (9.5%) | NA | NA | 23 / 55 (42%) | 3 / 11 (27%) | NA | NA |
| **NA** | 18 | 44 | 87 | 62 | 60 | 42 | 15 | 63 | 10 | 38 | 42 | 68 | 10 | 2 | 51 | 89 |
| **Hypomanic Episodes** | | | | | | | | | | | | | | | | |
| **Mean (SD)** | NA | NA | NA | 1 (0.4) | 4 (7) | NA | NA | NA | NA | 3 (3) | NA | NA | NA | NA | NA | NA |
| **NA** | 49 | 44 | 87 | 62 | 112 | 42 | 15 | 127 | 19 | 59 | 57 | 68 | 65 | 13 | 51 | 89 |
| **Manic Episodes** | | | | | | | | | | | | | | | | |
| **Mean (SD)** | NA | 8 (7) | 5 (6) | 3 (2) | 1 (2) | NA | 30 (25) | 49 (62) | NA | 4 (3) | NA | NA | NA | 4 (2) | NA | 5 (5) |
| **NA** | 49 | 23 | 75 | 21 | 3 | 42 | 11 | 56 | 19 | 40 | 57 | 68 | 65 | 6 | 51 | 66 |
| **Depressive Episodes** | | | | | | | | | | | | | | | | |
| **Mean (SD)** | NA | 9 (6) | 9 (11) | 3 (2) | 5 (10) | NA | 25 (28) | 41 (49) | NA | 4 (3) | NA | NA | NA | 5 (6) | NA | 8 (8) |
| **NA** | 49 | 23 | 76 | 21 | 1 | 42 | 11 | 56 | 19 | 40 | 57 | 68 | 65 | 4 | 51 | 67 |
| **Mixed Episodes** | | | | | | | | | | | | | | | | |
| **Mean (SD)** | NA | NA | NA | NA | NA | NA | NA | NA | NA | NA | NA | NA | NA | NA | NA | NA |
| **NA** | 49 | 44 | 87 | 62 | 112 | 42 | 15 | 127 | 19 | 59 | 57 | 68 | 65 | 13 | 51 | 89 |
| **Psychiatric Hospitalizations** | | | | | | | | | | | | | | | | |
| **Mean (SD)** | NA | NA | 1 (3) | NA | NA | 2 (3) | NA | NA | NA | NA | NA | NA | NA | NA | NA | 1 (2) |
| **NA** | 49 | 44 | 9 | 62 | 112 | 30 | 15 | 127 | 19 | 59 | 57 | 68 | 65 | 13 | 51 | 18 |

**Table S3. Inclusion and Exclusion Criteria for Each Cohort**

| Site | Inclusion | Exclusion |
|---|---|---|
| **Barcelona** | Age 18-65, right-handed, current IQ in the normal range (>70). All patients were diagnosed using DSM-IV and Research Diagnostic Criteria (RDC), based on a detailed clinical interview and review of case notes. | History of neurological disease or brain trauma, alcohol/substance abuse in the 12 months prior to participation, or IQ <70. Healthy controls met the same exclusion criteria as the patients, plus history of mental illness and/or treatment with psychotropic medication other than non-regular use of benzodiazepines or similar drugs for insomnia, or a first-degree relative with symptoms consistent with major psychiatric disorder and/or having received any form of in- or outpatient psychiatric care. |
| **Moodinflame** | Adult male and female subjects who were free of inflammation-related symptoms including fever and current or recent infectious or inflammatory disease, uncontrolled systemic disease, uncontrolled metabolic disease or other significant uncontrolled somatic disorders known to affect mood. They did not use somatic medication known to affect mood or the immune system, such as corticosteroids, non-steroid anti-inflammatory drugs and statins. Female candidates who were pregnant or recently gave birth were excluded. Patients were allowed to continue their regular psychopharmacological treatment. They were euthymic at the time of scanning as indicated by an Inventory of Depressive Symptoms - Clinician Version (IDS-C30) score o22 and a Young Mania Rating Scale (YMRS) score o 12, respectively. Patients with any other current primary major psychiatric diagnosis were excluded including: schizophrenia, schizoaffective disorder, anxiety disorder and substance use disorders. | HC did not have any current or lifetime psychiatric diagnosis. |
| **FOR 2017 Munster** | Inclusion criteria: age 17-65 years; patients were diagnosed of bipolar I disorder by SCID-Interview, currently depressed (HAMD >= 18); | Participants with neurological or general medical conditions, substance dependence, or a verbal IQ of 80 or lower were excluded. Healthy control subjects with current or past mental disorders based on the Structured Clinical Interview for DSM-IV-TR (SCID-I) or who had ever used psychotropic medication were also excluded. Any MRI contraindications. |
| **Pittsburgh** | Structured Clinical Interview for DSM-IV for Axis I Diagnoses. Pittsburgh Inclusion criteria: Euthymic individuals with BP I disorder (diagnosed according to the criteria of DSM-IV (APA, 1994) and Bipolar I section of the SCID (First et al., 1994): right-handed; HRSD-25 score of equal or less than 7 on the 17 item portion and YMRS less than or equal to 10; meets criteria for being euthymic; Individuals with BP disorder currently in depressed episode (diagnosed according to the criteria of DSM-IV and SCID criteria): right-handed; not currently in a manic, hypomanic or mixed episode; Healthy controls: Right-handed; HRSD-25 score of equal or less than 7 on the 17 item portion and a YMRS score of equal or less than 10; No first degree relatives with diagnosed psychiatric disorders. | History of head injury or neurological illness; Mini-Mental State Exame <24; IQ <85 assessed with NART; Visual disturbance (<20/40); current dependence on alcohol or drug use; contraindication for MRI; pregnancy; lack of English proficiency. |
| **Sydney** | Age 12-30. Diagnostic Interview for Genetic Studies (for 22-30 year-olds); Kiddie-SADS (for 12-21 year-olds) | Control participants were defined as those who did not have a first-degree relative with either BD I or II, recurrent major depressive disorder (MDD), schizoaffective disorder, recurrent substance abuse or any past psychiatric hospitalization. In the BD group only those with a diagnosis of BD I or II were included. |

| | | |
|---|---|---|
| **Milano** | Subjects were assessed for diagnosis by trained raters using the Schedules Clinical Assessment Neuropsychiatry. We defined patients with a positive history of psychotic features if they had at least one episode of depression or mania either with delusions or hallucinations. | Participants with neurological or general medical conditions, substance dependence, or a verbal IQ of 80 or lower were excluded. Healthy control subjects with current or past mental disorders based on the Structured Clinical Interview for DSM-IV-TR (SCID-I) or who had ever used psychotropic medication were also excluded. |
| **Yale/Olin_32 gradients** | Patients were identified through outpatient clinics and community mental health facilities in the Hartford area. Inclusion criteria for patients included: diagnosis of bipolar I disorder as determined by the Structured Clinical Interview [First et al., 2002], administered by experienced research clinicians, and IQ > 80. Unrelated healthy comparison subjects were recruited through media advertisements and flyers. In the interest of sample validity, history of major depressive episodes, comorbid anxiety disorders, and history of substance abuse/dependence were allowed for all subjects | Current substance or alcohol abuse/dependence; history of a major medical or neurological condition. |
| **Yale/Olin_55 gradients** | Patients were identified through outpatient clinics and community mental health facilities in the Hartford area. Inclusion criteria for patients included: diagnosis of bipolar I disorder as determined by the Structured Clinical Interview [First et al., 2002], administered by experienced research clinicians, and IQ > 80. Unrelated healthy comparison subjects were recruited through media advertisements and flyers. In the interest of sample validity, history of major depressive episodes, comorbid anxiety disorders, and history of substance abuse/dependence were allowed for all subjects | Current substance or alcohol abuse/dependence; history of a major medical or neurological condition. |
| **Capetown** | All recruited participants were aged between 19-40 years. Patients were currently stable outpatients, euthymic, with SCID-I confirmed DSM-IV diagnoses of BP. Healthy controls were screened out against SCID-I psychiatric disorders (First et al. 2002). | Participants were excluded if they reported head injury, self/family history of epilepsy, or were receiving pharmaceutical treatment for a general medical condition. |
| **NUIG-CHRM2** | ages 18–65, were recruited by referral or public advertisement from the western regions of Ireland's Health Services. A diagnosis of BD was confirmed using the Diagnostic and Statistical Manual of Mental Disorders, 4th edition (DSM-IV-TR) Structured Clinical Interview for DSM Disorders (American Psychiatric Association, 2013) conducted by an experienced psychiatrist. Mood and anxiety symptom severity were assessed using the Hamilton Depression Rating Scale (HDRS-21), Hamilton Anxiety Rating Scale (HARS), and Young Mania Rating Scale (YMRS) at MRI scanning, and in BD a diagnosis of euthymia was defined by HDRS <8, YMRS <7, and HARS <18 scores. | Exclusion criteria included neurological disorders, learning disability, comorbid misuse of substances/alcohol, other Axis-1 disorders, history of head injury resulting in loss of consciousness for >5 min, along with a history of oral steroid use in the previous 3 months. Healthy controls had no personal history of a psychiatric illness or history among first-degree relatives, defined using the Structured Clinical Interview for DSM-IV Non-patient edition (American Psychiatric Association, 1994). |
| **Creteil** | Inclusion criteria for study participation were ages between 18 and 65, no history of alcohol or drug abuse/dependence, no history of mental retardation, no previous head trauma with loss of consciousness, and no current or past cardiac or neurological disease. Euthymic individuals with BP I disorder (diagnosed according to the criteria of DSM-IV (APA, 1994) and Bipolar I section of the SCID (First et al., 1994): right-handed; HRSD-25 score of equal or less than 7 on the 17 item portion and YMRS less than or | Exclusion criteria for all participants consisted of a history of neurological disease or head trauma with loss of consciousness and contraindications for magnetic resonance imaging (MRI). We excluded subjects with any significant cerebral anatomic anomaly. |

| | | |
|---|---|---|
| | equal to 10; meets criteria for being euthymic; Individuals with BP disorder currently in depressed episode (diagnosed according to the criteria of DSM-IV and SCID criteria): right-handed; not currently in a manic, hypomanic or mixed episode; Healthy controls: no previous psychiatric history; right-handed; HRSD-25 score of equal or less than 7 on the 17 item portion and a YMRS score of equal or less than 10; No first degree relatives with diagnosed psychiatric disorders | |
| **Deakin** | Patients were screened with the Structured Clinical Interview for DSMIV-TR by trained clinicians and were diagnosed with mania (bipolar I disorder, schizoaffective disorder, or a substance-induced mood disorder). Individuals presenting with an acute manic episode with psychotic features and who had not been previously treated for a manic episode were stabilized on a combination of quetiapine plus lithium in an open label manner as part of a routine care protocol. Following provision of informed oral and written consent, patients were randomized after remission (2–3 months), to lithium or quetiapine monotherapy. Patients were required to have been on a combination of quetiapine and lithium as standard therapy for at least 1 month prior to randomization, and to have a score of at least 20 on the Young Mania Rating Scale (YMRS)33 during their acute manic episode. Female patients were required to be using effective contraception if they were sexually active and of childbearing age. Patients of 15–25 years of age, who were fluent in English, had the capacity to provide informed consent, and comply with study procedures, were randomized to treatment allocation of either lithium or quetiapine monotherapy. | Patients were excluded for known or suspected clinically relevant systemic medical disorder, organic mental disease or a history of epilepsy; sensitivity or allergy to quetiapine, lithium or any additives in the medication prior use of medication with a cytochrome P450 3A4 inhibiting effect in the 14 days preceding enrolment; inability to comply with the requirements of giving informed consent or the treatment protocol; immediate risk of self-harm or of harming others; pregnancy or breastfeeding; or diabetes mellitus. |
| **NUIG-GBS** | BD type I was confirmed using the Diagnostic and Statistical Manual of Mental Disorders, 4th edition (DSM-IV) Structured Clinical Interview for DSM Disorders (American Psychiatric Association, 1994). | Exclusion criteria for all participants included a history of medical or neurological illness, history of head injury resulting in loss of consciousness for over 5 min, history of substance abuse in the past year, learning disability, and oral steroid use in previous 3 months. Further exclusion criteria for controls included personal or family history of psychotic or affective disorder in first- or second-degree relatives. Additional patient exclusion criteria included a lifetime co-morbid DSM-IV Axis I disorder. |
| **Grenoble** | BD (euthymic) 18-65 | no history of alcohol or drug abuse; current or past neurological and/or medical diseases affecting cognition; history of head trauma with loss of consciousness; metal implants. Additionally, we excluded (a) for BD, any current other Axis I psychiatric disorder and sismotherapy during the precedent year; (b) for HC, past or present psychiatric disorder and family history of psychiatric disorders, as well as any medical treatment affecting cerebral activity |
| **Mannheim** | Age 18-65. Euthymia was defined by a Hamilton Depression Rating Scale (HAM-D) (19) score <5 and a Young Mania Rating Scale (20) score <8 for at least 8 weeks. Inclusion criteria for study participation comprised no history of alcohol or drug abuse/dependence, no history of mental retardation, and no current or past cardiac or neurological disease. Healthy comparison subjects were free of any past or present psychiatric | Current substance or alcohol abuse/dependence; history of a major medical or neurological condition. |

| | | |
|---|---|---|
| | disorder (as assessed by the Mini-International Neuropsychiatric Interview | |
| **FOR-2017 Marburg** | Age 18-65 | Participants with neurological or general medical conditions, substance dependence, or a verbal IQ of 80 or lower were excluded. Healthy control subjects with current or past mental disorders based on the Structured Clinical Interview for DSM-IV-TR (SCID-I) or who had ever used psychotropic medication were also excluded. |

**Table S4. MRI data Acquisition Parameters for Each Cohort**

| Site | T1-weighted scan | Diffusion-weighted scan |
|---|---|---|
| **Barcelona** | 3D T1-weighted enhanced fast gradient echo (EFGRE3D) 1.5T GE Signa, 180 axial slices, no gap, voxel size 0.47×0.47×1mm3, TI 710ms, TE 3.93ms, RE 2000ms, flip angle 15 | GE Signa 1.5T, voxel size 2.2 × 2.2 × 3 mm3, 3 slice thickness, 55 directions, 1500 b-value s/mm2, 1 b=0 scan |
| **Moodinflame** | One high-resolution (1 mm isotropic) T1-weighted image for each participant (SPGR, TR¼9758 ms, TE¼4.59 ms, ip angle 8°, FOV¼ 220 220 mm, voxel size 0.859 0.859, slice thickness 1.2 mm) covering the entire cerebrum. The total acquisition time for this sequence was less than 5 min. | Diffusion-weighted (DW) data were acquired using a single-shot pulsed gradient spin echo EPI sequence (TR¼8041 ms, TE¼59.98 ms) on a 3-T MR scanner (Intera, Philips Medical Systems, Best, the Netherlands) equipped with an eight-channel head coil. For each subject, 32 DW images were collected along non-collinear directions, with a maximum gradient strength of 40 mT/m and a b-value of 1000 s/mm2. In addition, one non-DW volume (b E0, referred to in the following as a b0 image) was acquired. Each volume consisted of 55 transverse slices (FOV¼240 240 mm, voxel size¼ 2.5 mm isotropic, no gap). The total acquisition time for this sequence was less than 6 min. |
| **FOR-2017 Munster** | T1-weighted images were acquired using a 3 T MRI scanner (Tim Trio, Siemens, Erlangen, Germany; Münster: Prisma, Siemens, Erlangen, Germany), a 20-channel head matrix Rx-coil was used. Following a quality assurance protocol, based on regular measurements of an MRI phantom [60], whole-brain T1-weighted scans were obtained using a fast gradient echo MP-RAGE sequence with a slice thickness of 1.0 mm, 176 (at Münster site: 192) sagittal orientated slices and FOV of 256 mm with the following parameters: Marburg: TR = 1.9 s, TE = 2.26 ms, TI = 900 ms, flip angle = 8°; Münster: TR = 2.13 s, TE = 2.28m s, TI = 900 ms, flip angle = 9°. | DTI scans were acquired using an epi2d sequence (TR 7300 ms, TE 90 ms, FOV 320 mm, phase encoding anterior-posterior, 56 slices with 2.5 mm slice thickness) with a final voxel resolution of 2.5 × 2.5 × 2.5 mm3. For all patients, two sets of 30 diffusion-weighted images (b = 1000 s/mm2) and four non-diffusion weighted images (b = 0 s/mm2) were acquired. |
| **Pittsburgh** | 3D T1-weighted magnetization prepared rapid acquisition gradient echo (MPRAGE); 3T Siemens Trio. Axial acquisition, 192 slices, 0mm slice gap, 1x1x1 mm3, TI = 900ms, TE = 3.29ms, TR = 2200ms, flip angle = 9 | Siemens TrioTrim 3T, 1 acquisition,2 × 2 mm3 voxel size, 2 slice thickness, 41 directions, 1000 b-value<br>s/mm2, 1 b=0 scan |
| **Sydney** | T1-weighted images were acquired using a 3T Philips Achieva scanner (Royal Philips Electronics, Amsterdam, The Netherlands) located at Neuroscience Research Australia in Sydney. One-hundred-and-eighty sagittal T1-weighted three-dimensional turbo field-echo images were obtained (TR/TE – 5.5/2.5ms, flip angle – 8°, field of view – 256 x 256 x 180mm3, voxel size – 1 x 1 x 1mm, scan time – 371s), preceded by a one-minute standard scout image for head positioning and resolution of sensitivity variations. | Diffusion MRI data were acquired with a 3-T Philips Achieva scanner with an 8-channel head coil. 32 directional DTI data (b =1000 s/mm2, with one non-diffusion-weighed image) was acquired using a single-shot echo planar imaging (EPI) sequence. The imaging parameters were as follows: TR = 7767 ms, TE = 68 ms, 55 slices, slice thickness = 2.5 mm, gap = 0 mm, acquisition matrix size = 96 × 96 (field of view = 240×240×137.5 mm), flip angle = 90°, reconstructed to yield 1 mm × 1 mm × 2.5 mm voxels (where the longer dimension is along the dorsoventral axis). |

| | | |
|---|---|---|
| **Milano** | 3T Siemens Allegra (echo time = 3.93ms, repetition time = 2300 ms, image size = 256×256, slice thickness 1mm | Philips Intera 3T, 1 acquisition, 2.14x3.71 mm3 voxel size, 2.3 mm slice thickness, 35 directions, 900 b-value s/mm2, 1 b=0 scan. |
| **Yale/Olin_32 gradients** | Structural images were acquired using a T1-weighted, 3D magnetization-prepared rapid gradient-echo (MPRAGE) sequence (TR/TE/TI = 2200 / 4.13 / 766 ms. flip angle = 13, voxel size [isotropic] = 0.8 mm, image size = 240 x 320 x 280 voxels), with axial slices parallel to AC-PC line. To increase signal to noise ratio, four volumes were acquired per subject. | Diffusion-weighted MR images were acquired using single-shot echo planar imaging (TR/TE=6300/81 ms, field of view = 22 3 22 cm, acquisition matrix = 128x3x128, voxel size = 1.7 3x1.73x3.0mm) with a twice-refocusing spin echo sequence to minimize eddy-current induced distortion. The sequence consisted of 32 non-collinear diffusion weighted directions, one diffusion weighting values (1000s/mm2) along with a single non-diffusion-weighted image (b= 0). |
| **Yale/Olin_55 gradients** | Structural images were acquired using a T1-weighted, 3D magnetization-prepared rapid gradient-echo (MPRAGE) sequence (TR/TE/TI = 2200 / 4.13 / 766 ms. flip angle = 13, voxel size [isotropic] = 0.8 mm, image size = 240 x 320 x 280 voxels), with axial slices parallel to AC-PC line. To increase signal to noise ratio, four volumes were acquired per subject. | Diffusion-weighted MR images were acquired using single-shot echo planar imaging (TR/TE5=6300/81 ms, field of view = 22 3 22 cm, acquisition matrix= 128x3x128, voxel size= 1.73x1.73x3.0mm) with a twice-refocusing spin echo sequence to minimize eddy-current induced distortion. The sequence consisted of 55 non-collinear diffusion weighted directions, two diffusion weighting values (b=0 and 800s/mm2) and three non-diffusion weighted (b=0) images. The acquisition lasted 6.2 minutes and contained 58 sets of images, with 45 contiguous, interleaved axial slices per volume (slice thickness 5 3.0mm) covering the whole brain. |
| **CIAM Cape Town** | Structural MPRAGE (van der Kouwe et al., 2008) sequence :TR = 2530 ms, graded TE = 1.53, 3.21, 4.89, 6.57 ms, flip angle = 7°, FOV = 256 mm, slice thickness = 1.33 mm, 128 slices, voxel size 1.3x1.0x1.3, scan time 8:06. Single channel coil used within a 3T Siemens Allega head scanner. | Diffusion-weighted MR images were acquired within the same hour as the structural MRI within 3T Allegra Siemens head scanner, two acquisitions were collected (1 x A-P, 1 x P-A), TR 7600ms, TE 88ms, FOV 230mm, Voxel size 1.8x1.8x2.0mm, 55 slices with slice thickness of 2mm, scan time 4:18 for each direction |
| **NUIG-CHRM2** | High-resolution three-dimensional T1-weighted turbo field echo magnetization-prepared rapid gradient-echo (MPRAGE) sequence was acquired using an eight-channel head coil (parameters: repetition time [TR]/echo time [TE]= 8.5/3.046 ms, 1 mm3 isotropic voxel size). | Diffusion-weighted images were acquired at b= 1200 s/mm2 along with a single nondiffusion-weighted image (b= 0), using high-angular resolution diffusion imaging (HARDI) involving 61 diffusion gradient directions, 1.8 · 1.8 · 1.9 mm voxel dimension, and field of view 198 · 259 · 125 mm. |
| **Creteil** | high-resolution T1-weighted acquisition (echo time, 2.98 milliseconds; repetition time, 2300 milliseconds; 160 sections; voxel size, 1.0 × 1.0 × 1.1 or 1.0 mm) | DW sequence along 41 directions (voxel size, 2.0 × 2.0 × 2.0 mm; b = 1000 s/mm2 plus 1 image in which b = 0; echo time, 87 or 84 milliseconds; repetition time, 14 000 milliseconds; 60 or 64 axial sections) |
| **Deakin** | Siemens 3T TrioTim, TR/TE/TI=2000/2.24/900ms, Flip Angle =9, 232x256 matrix, 192 slices saggital, 0.9mm isotropic | TR=8800ms, TE=99ms, Flip angle=90, 1024x1024 matrix, 64 slices, 2mm slice thickness with 2mm slice gap, 10 b0, 58 directions. b0=2000 s/mm^2 |
| **NUIG-GBS** | T1-weighted magnetization prepared acquisition of gradient echo (MPRAGE) sequence was acquired with the imaging parameters: repetition time 1140 ms; echo time 4.38 ms; inversion time 600 ms; flip angle 15; matrix size 256 × 256; an in-plane pixel size 0.9 × 0.9 mm2; slice thickness of 0.9 mm. | Diffusion MRI data were acquired using an eight-channel head coil with an echo planar image diffusion sequence acquired with parallel imaging, 64 optimized diffusion gradient directions with b = 1300 s/mm2, seven non-diffusion weighted images, repetition time = 8100 ms, echo time = 95 ms, field of view = 240 × 240 mm2, matrix = 96 × 96, in-plane voxel size of 2.5 × 2.5 mm2, slice thickness = 2.5 mm, 60 slices. |
| **Grenoble** | T1-weighted high-resolution 3D volume (resolution: 0.8 × 0.8 × 0.8 mm) | Philips Achieva 3T, 1 acquisition, 30 directions; 3 x 3 mm3 voxel size, 3-mm-thick axial slices; b value = 1,400 s/mm2) |

| | | |
|---|---|---|
| **Mannheim** | One high-resolution T1-weighted (1.5 T GE Signa scanner), three-dimensional magnetic resonance imaging (MRI) sequence (slice thickness=1.3 mm; field of view=24 cm; 256×256×128 matrix) | Siemens TrioTrim 3T, 1 acquisition,2 × 2 mm3 voxel size, 41gradients, 1000 b-value, 1 b=0 scan |
| **FOR-2017 Marburg** | T1-weighted images were acquired using a 3 T MRI scanner (Tim Trio, Siemens, Erlangen, Germany; Münster: Prisma, Siemens, Erlangen, Germany), a 12-channel head matrix Rx-coil was used. Following a quality assurance protocol, based on regular measurements of an MRI phantom [60], whole-brain T1-weighted scans were obtained using a fast gradient echo MP-RAGE sequence with a slice thickness of 1.0 mm, 176 (at Münster site: 192) sagittal orientated slices and FOV of 256 mm with the following parameters: Marburg: TR = 1.9 s, TE = 2.26 ms, TI = 900 ms, flip angle = 8°; Münster: TR = 2.13 s, TE = 2.28m s, TI = 900 ms, flip angle = 9°. | DTI scans were acquired using an epi2d sequence (TR 7300 ms, TE 90 ms, FOV 320 mm, phase encoding anterior-posterior, 56 slices with 3 mm thickness in Marburg) with a final voxel resolution of 2.5 × 2.5 × 2.5 mm3. For all patients, two sets of 30 diffusion-weighted images (b = 1000 s/mm2) and four non-diffusion weighted images (b = 0 s/mm2) were acquired. |

***Whole-brain measures derived from the connectome***

Connectome-derived graph theory metrics offer a quantitative framework for characterizing the topological organization of brain networks. These metrics are commonly grouped into three domains: integration, segregation, and influence (Rubinov & Sporns, 2010; Sporns, 2011). In this study, we computed graph theory measures from structural connectivity matrices derived using diffusion-weighted MRI tractography. To provide a comprehensive assessment of network architecture, metrics were calculated on both weighted and unweighted versions of the connectome. In unweighted networks, only the presence or absence of edges is considered (i.e., binary connections). This approach highlights topological properties and simplifies comparisons across individuals, especially when the reliability of edge weights may be uncertain. In weighted networks, edges carry quantitative values that reflect the strength or microstructural organization of the white matter connections. We used two weighting schemes; NOS: the number of reconstructed streamlines between each pair of brain regions, reflecting macrostructural connectivity. FA: the mean FA value along the streamlines connecting two regions, indexing microstructural organization of the white matter tracts. Both metrics offer complementary information: streamline count captures the quantity of fiber pathways, while FA reflects the quality or coherence of these tracts. Analyzing both enables a richer understanding of how macrostructural features relate to clinical outcomes in BD. Calculating both forms allows for cross-validation of findings and addresses potential methodological biases. While weighted networks are more biologically informative, they can also be more sensitive to noise and thresholding. Using both types ensures a more robust and nuanced understanding of brain network organization and its relationship to clinical variables in BD. Table S4 provides definitions and formulas for each graph metric, including descriptions of how they differ between weighted and unweighted implementations.

| ***Metric Name*** | ***Formula*** | ***Meaning*** |
|---|---|---|
| Network Density ($D$) | $D = \frac{2E}{N(N-1)}$<br>● *E* is the number of edges in the network,<br>● *N* is the number of nodes. | Proportion of existing connections out of all possible connections in the network, $\frac{N(N-1)}{2}$. Reflects how densely connected the graph is. Only meaningful in unweighted networks. |
| Degree ($k_i$ ) | $k_i = \Sigma_j a_{ij}$<br>● *Unweighted:* $a_{ij}$ = 1 if edge exists, 0 otherwise<br>● *Weighted analog:* Strength, $s_i = \Sigma_j w_{ij}$<br>● $s_i$ is the strength of node *i*<br>● $w_{ij}$ is the weight of the edge between node *i* and node *j*<br>● *N* is the set of all nodes | Measures a node's direct connectivity. Nodes with higher strength (or degree centrality) are often more active or important in communication. In unweighted networks, degree is the count of edges. In weighted networks, the corresponding measure is strength, summing all edge weights. |
| Clustering Coefficient ($C_i$ ) | $C_i = \frac{2T}{k_i(k_i-1)}$<br>● $C_i$ is the clustering coefficient of node ii<br>● $T_i$ is the number of triangles (i.e., closed triplets) through node *i*. - Weighted: $T_i$ includes geometric mean of edge weights<br>● $K_i$ is the degree of node ii, or the number of connections it has | Reflects the tendency of a node's neighbors to be interconnected. A higher $C_i$ indicates a greater tendency for a node's neighbors to also be connected, reflecting localized connectivity or community structure, and implies strong local segregation. In weighted graphs, the strength of connections is considered in triangle formation. |

| | | |
|---|---|---|
| Characteristic Path Length ($L$) | $$L = \frac{1}{N(N-1)} \Sigma_{i \neq j} d_{ij}$$ ● *N* is the total number of nodes in the network<br>● *dij* is the shortest path length between node *i* and node *j* | Measures the average shortest path between all pairs of nodes. Shorter paths imply more efficient communication. In weighted networks (e.g., streamline count or FA), edge weights are interpreted as connection strength, and path lengths are computed as inverses of weights. |
| Local Efficiency ($E_{loc}$) | $$E_{loc,i} = \frac{1}{k_i(k_i - 1)} \Sigma_{j,h \in N_i} \frac{1}{d_{jh}}$$ ● $T_i$ is the set of neighbors of node i<br>● $K_i$ is the degree of node i<br>● $d_{jh}^{(i)}$ is the shortest path length between neighbors *j* and *h* within the subgraph composed of *i*'s neighbors (excluding node *i*) | Measures local fault tolerance, or how efficiently a node's neighbors communicate when the node is removed. Higher $E_{loc}$ suggests robust local connectivity that can preserve communication even if a central node is disrupted. Can be computed using weighted or unweighted paths within each node's subgraph. |
| Global Efficiency ($E_{glob}$) | $$E_{glob} = \frac{1}{N(N-1)} \Sigma_{i \neq j} \frac{1}{d_{ij}}$$ ● *N* is the number of nodes<br>● *dij* is the shortest path length between nodes *i* and *j* | Reflects the efficiency of parallel information transfer across the network. Higher values indicate better global communication. $E_{glob}$ increases when nodes are more directly connected, supporting faster and more efficient communication. In weighted networks, the inverse of edge weights is used. |
| Betweenness Centrality ($BC_i$) | $$BC_i = \Sigma_{s \neq i \neq t} \frac{\sigma_{st}(i)}{\sigma_{st}}$$ ● $BC(i)$ is the betweenness centrality of node *i*<br>● $\sigma_{st}$ is the total number of shortest paths from node ss to node tt<br>● $\sigma_{st}(i)$ is the number of those paths that pass through node ii<br>● *Weighted:* paths computed with inverse edge weights | Quantifies how often a node lies on the shortest path between other node pairs. High $BC(i)$ indicates that node ii is critical for efficient communication between many parts of the network, and implies a central role in network integration<br>In weighted graphs, paths account for connection strength. |

**Table S5. Connectome derived graph theory metrics.** Table lists each metric, its formula, and a brief interpretation. Unweighted (binary) calculations use edge presence only; weighted versions use streamline count (NOS) and fractional anisotropy (FA).

| Traditional Labels | NbN Categories | Medications |
|---|---|---|
| **Antipsychotics** | **NbN 1** - Primarily Dopamine Receptor Antagonist | Haloperidol, Zuclopenthizol, Perphenazine, Fluphenazine, Sulpiride, Amisulpride |
| | **NbN 2** - Dopamine and Other (Ser-nor) Monoamine Receptor Antagonists | Flupenthixol, Levomepromazine, Chlorpromazine, Trifluoperazine, Pipotiazine, Loxapine, Thioridazine, Olanzapine, Lurasidone, Ziprasidone, Risperidone, Paliperidone, Asenapine, Quetiapine, Clozapine |
| | **NbN 3** - Dopamine, Serotonin Receptor Partial Agonist/Antagonist | Aripiprazole |
| **Antidepressants** | **NbN 4** - Targeting Serotonin (Reuptake Inhibitors Multimodal) | Escitalopram, Citalopram, Paroxetine, Fluoxetine, Fluvoxamine, Sertraline, Trazodone |
| | **NbN 5** - Targeting Serotonin and Other Monoamines with different MOA | Mirtazapine, Venlafaxine, Duloxetine, Clomipramine, Dosulepin, Imipramine, Bupropion, Amitryptiline, Nortrptyline, Tranylcypromine, Moclobemide, Reboxetine, Agomelatine |
| **Antiepileptics** | **NbN 6** - Glutamate, Sodium, Calcium Channel Blockers | Carbamazepine, Oxcarbazepine, Lamotrigine, Pregabalin, Gabapentin |
| **Mood Stabilizer (Lithium)** | **NbN 7** - Lithium | Lithium |
| **Mood Stabilizer (Valproate)** | **NbN 8** - Valproate | Sodium Valproate |
| **Other Psychotropics** | **NbN 9** - Other | Benzodiazepine, Topiramate, etc |

**Table S6. Neuroscience Based Nomenclature.** This table summarizes the categorization of medications based on their underlying neurobiological and pharmacological profiles, following the Neuroscience-Based Nomenclature (NbN) Guidelines (Uchida et al., 2016). Medications were classified according to their primary mechanisms of action and neurobiological targets. Traditional medications were further categorized to provide a detailed understanding of their mechanisms. The table aims to offer clarity on how each medication aligns with specific neurobiological profiles and their therapeutic implications.

| *Model Family* | *Model Type* | *Equation (fixed-effects part)* | *Random term* | *Purpose* |
|---|---|---|---|---|
| **Diagnosis** | Main | WM = $\beta_0$+ $\beta_1$*Diagnosis + $\beta_2$*Age + $\beta_3$*Sex | (1 \| Site) | Test BD *versus* controls differences on structural connectivity measures |
| **Medications** | Main | WM = $\beta_0$+ $\beta_1$*$Med_1$ + $\beta_2$*Age + $\beta_3$*Sex | (1 \| Site) | Association of a single medication class on structural connectivity measure |
| | *Post-hoc* | WM = $\beta_0$+ $\beta_1$*$Med_1$ + $\beta_2$*Age + $\beta_3$*Sex + $\beta_4$*$Med_2$ + $\beta_5$*$Med_3$ + $\beta_6$*$Med_4$ + $\beta_7$*$Med_{5...}$ | (1 \| Site) | Class-specific associations with structural connectivity measures while adjusting for concurrent use of the other classes (polypharmacy) |
| | *Post-hoc* | WM = $\beta_0$+ $\beta_1$*$Med_1$ + $\beta_2$*Age + $\beta_3$*Sex + $\beta_4$*ManicEp + $\beta_5$*DepEp | (1 \| Site) | Class-specific associations with structural connectivity measures additionally adjusting for illness-course severity measures |
| **Illness course** | Main | WM = $\beta_0$+ $\beta_1$*IllnessSeverity + $\beta_2$*Age + $\beta_3$*Sex | (1 \| Site) | Associations of illness severity index with structural connectivity measures |
| | *Post-hoc* | WM = $\beta_0$+ $\beta_1$*IllnessSeverity + $\beta_2$*Age + $\beta_3$*Sex + $\beta_4$*$Med_1$+ $\beta_5$*$Med_2$+ $\beta_6$*$Med_3$+ $\beta_7$*$Med_4$+$\beta_8$*$Med_5$ ... | (1 \| Site) | Associations of illness severity index with structural connectivity measures while controlling for all medication classes (polypharmacy) |

**Table S7. Summary of linear mixed-effects models (LMMs) used to assess associations between white-matter metrics and diagnosis, medication use, and illness severity**. We used linear mixed models (LMMs) to assess the relationship between whole-brain normally distributed measures and diagnosis, while adjusting for age, sex and site. In this model, diagnosis, age, and sex were included as fixed effects, and site was included as a random intercept to account for between-site variability (R v4.2.1). *Post-hoc* models account for polypharmacy and illness course to isolate specific effects.

| | |
|---|---|
| **BD Type**, n / N (%) | |
| BD I | 201 / 243 (83%) |
| BD II | 42 / 243 (17%) |
| NR | 207 |
| **Mood Phase**, n / N (%) | |
| Depressed | 36 / 359 (10%) |
| Euthymic | 308 / 359 (86%) |
| Hypomanic | 4 / 359 (1.1%) |
| Manic | 9 / 359 (2.5%) |
| Mixed | 2 / 359 (0.6%) |
| NR | 91 |
| **Duration of Illness** (Years) | |
| Mean (SD) | 16 (11) |
| Range | 0.6-45 |
| Median, | 14 |
| NR | 65 |
| **Age of Onset** (Years) | |
| Mean (SD) | 21 (9) |
| Median | 19 |
| Early, n, %/N | 148, 38% |
| Intermediate, n, %/N | 199, 52% |
| Late, n, %/N | 39, 10% |
| NR | 64 |
| **History of Psychosi**s, n / N (%) | |
| 0 | 112 / 228 (49%) |
| 1 | 116 / 228 (51%) |
| NR | 222 |
| **Hypomanic Episodes** | |
| Mean (SD) | 3 (4) |
| NR | 378 |
| **Manic Episodes** | |
| Mean (SD) [N] | 6 (9) [N =282] |
| Range | 0-60 |
| Median | 3 |
| NR | 228 |
| **Depressive Episodes** | |
| Mean (SD) [N] | 9 (12) [N=286] |
| Range | 0-72 |
| Median | 5 |
| NR | 221 |
| **Mixed Episodes** | |
| n / N (%) | 1 / 450 (0.2%) |
| NR | 449 |
| **Psychiatric Hospitalizations** | |
| Mean (SD) | 1 (3) |
| NR | 402 |

**Table S8. Clinical and Sociodemographic Details of Bipolar disorder participants.** BD = Bipolar Disorder; n/N = Sample/Total Sample Size; NR = Not Reported; SD = Standard Deviation. Age of Onset: Early: < 18 years ; 18 years < intermediate < 35 years; Late: > 35 years

| | (i) Connectivity strength (mean ± SD) | | (ii) Percentage change in BD (%) vs. controls | (iii) The magnitude of network component effect (t-value) |
|---|---|---|---|---|
| Suprathreshold connections | HC, N = 509 | BD, N = 450 | | |
| Left Cerebellum Cortex to Left Thalamus Proper | 0.346 ± 0.099 | 0.342 ± 0.094 | ↓ 1.16 | 3.72 |
| Left Thalamus Proper to Left Caudate | 0.331 ± 0.055 | 0.338 ± 0.044 | ↑ 2.11 | 3.27 |
| Left Thalamus Proper to Left Putamen | 0.388 ± 0.059 | 0.398 ± 0.062 | ↑ 2.58 | 3.58 |
| Left Putamen to Left Pallidum | 0.375 ± 0.049 | 0.370 ± 0.048 | ↓ 1.33 | 2.8 |
| Left Putamen to Left Ventral DC | 0.385 ± 0.067 | 0.382 ± 0.064 | ↓ 0.78 | 4.44 |
| Left Pallidum to Left Ventral DC | 0.388 ± 0.074 | 0.384 ± 0.067 | ↓ 1.03 | 6.26 |
| Left Hippocampus to Left Ventral DC | 0.317 ± 0.046 | 0.323 ± 0.043 | ↑ 1.89 | 4.05 |
| Left Amygdala to Left Ventral DC | 0.279 ± 0.056 | 0.287 ± 0.046 | ↑ 2.87 | 2.98 |
| Left Cerebellum Cortex to Right Cerebellum Cortex | 0.246 ± 0.071 | 0.229 ± 0.069 | ↓ 6.91 | 2.42 |
| Left Thalamus Proper to Right Thalamus Proper | 0.316 ± 0.075 | 0.334 ± 0.060 | ↑ 5.70 | 7.91 |
| Right Thalamus Proper to Right Caudate | 0.308 ± 0.063 | 0.315 ± 0.046 | ↑ 2.27 | 3.86 |
| Right Thalamus Proper to Right Putamen | 0.369 ± 0.054 | 0.378 ± 0.059 | ↑ 2.44 | 17.23 |
| Right Caudate to Right Putamen | 0.340 ± 0.054 | 0.354 ± 0.042 | ↑ 4.12 | 5.63 |
| Right Thalamus Proper to Right Pallidum | 0.371 ± 0.068 | 0.389 ± 0.053 | ↑ 4.85 | 5.23 |
| Right Caudate to Right Pallidum | 0.367 ± 0.053 | 0.372 ± 0.051 | ↑ 1.36 | 4.94 |
| Right Putamen to Right Pallidum | 0.371 ± 0.057 | 0.376 ± 0.046 | ↑ 1.35 | 2.18 |
| Right Thalamus Proper to Right Hippocampus | 0.332 ± 0.071 | 0.342 ± 0.064 | ↑ 3.01 | 9.85 |
| Right Caudate to Right Hippocampus | 0.330 ± 0.071 | 0.321 ± 0.074 | ↓ 2.73 | 2.1 |
| Right Hippocampus to Right Amygdala | 0.300 ± 0.080 | 0.272 ± 0.078 | ↓ 9.33 | 5.43 |
| Right Caudate to Right Accumbens area | 0.295 ± 0.062 | 0.306 ± 0.056 | ↑ 3.73 | 4.43 |
| Right Putamen to Right Accumbens area | 0.303 ± 0.058 | 0.302 ± 0.051 | ↓ 0.33 | 1.75 |
| Right Cerebellum Cortex to Right Ventral DC | 0.340 ± 0.095 | 0.322 ± 0.094 | ↓ 5.29 | 1.82 |
| Right Thalamus Proper to Right Ventral DC | 0.381 ± 0.072 | 0.390 ± 0.053 | ↑ 2.36 | 7.99 |
| Right Caudate to Right Ventral DC | 0.299 ± 0.067 | 0.305 ± 0.065 | ↑ 2.01 | 4.07 |
| Right Amygdala to Right Ventral DC | 0.293 ± 0.065 | 0.301 ± 0.060 | ↑ 2.73 | 11.93 |
| Left Hippocampus to ctx lh fusiform | 0.268 ± 0.061 | 0.271 ± 0.061 | ↑ 1.12 | 3.92 |
| Left Pallidum to ctx lh superiorfrontal | 0.392 ± 0.054 | 0.385 ± 0.056 | ↓ 1.79 | 1.74 |

**Table S9. Fractional Anisotropy-weighted subnetwork graph component showing dysconnectivity in bipolar disorder relative to controls.** Table shows the set of connections comprised in the subnetwork graph component found to show a significant effect for the main effect of diagnosis. NBS ‘F-test’ covarying for age and gender showing a dysconnected component in the bipolar group relative to controls (t>1.5, $p_{FWE}$= .0002, d=.2). (i) Connectivity strength FA for both groups and relative (ii) percentage decrease in strength for BD compared with controls; (iii) the magnitude of subnetwork component difference (t-value). This network showed 0.33–9.3% reduced connection strength associated with FA-edge weighting, and 1.12-5.70% increased connection strength in BD versus controls; components of the network ranged in magnitude of difference from a T>1.74 to 17.23, with the highest effects within and between basal ganglia and limbic connections. BD, bipolar disorder; HC, healthy controls; NBS, network-based statistics; SD, standard deviation; ventral DC, ventral diencephalon; ctx, cortical.

| Suprathreshold connections | (i) Connectivity strength (mean ± SD) HC, N = 509 | BD, N = 450 | (ii) Percentage change in BD (%) vs. controls | (iii) The magnitude of network component effect (t-value) |
|---|---|---|---|---|
| Left Caudate to Left Pallidum | 129.761 ± 96.149 | 165.949 ± 179.992 | ↑ 27.89 | 15.6 |
| Right Cerebellum Cortex to Right Thalamus Proper | 1,760.664 ± 2,592.723 | 739.787 ± 1,531.600 | ↓ 57.98 | 14.11 |
| Left Caudate to Right Caudate | 199.445 ± 264.590 | 254.854 ± 274.466 | ↑ 27.78 | 10.35 |
| Left Putamen to Right Putamen | 111.877 ± 159.945 | 61.766 ± 75.056 | ↓ 44.79 | 16.58 |
| Right Caudate to Right Putamen | 357.319 ± 259.943 | 401.762 ± 274.854 | ↑ 12.44 | 23.39 |
| Left Putamen to Right Pallidum | 90.523 ± 141.025 | 35.349 ± 53.212 | ↓ 60.95 | 14.6 |
| Right Caudate to Right Pallidum | 111.832 ± 95.389 | 172.846 ± 206.507 | ↑ 54.56 | 33.39 |
| Right Putamen to Right Pallidum | 698.623 ± 456.535 | 650.523 ± 410.754 | ↓ 6.88 | 10.24 |
| Right Pallidum to Right Hippocampus | 325.745 ± 448.025 | 174.762 ± 284.696 | ↓ 46.35 | 13.72 |
| Right Thalamus Proper to Right Amygdala | 163.664 ± 132.742 | 128.349 ± 123.732 | ↓ 21.58 | 11.16 |
| Right Pallidum to Right Amygdala | 105.662 ± 137.564 | 47.528 ± 69.415 | ↓ 55.02 | 15.02 |
| Right Amygdala to Right Accumbens area | 166.574 ± 216.997 | 84.578 ± 142.105 | ↓ 49.22 | 13 |
| Right Putamen to Right VentralDC | 424.000 ± 386.426 | 377.871 ± 310.114 | ↓ 10.88 | 21.9 |
| Right Pallidum to Right VentralDC | 509.665 ± 562.285 | 411.322 ± 467.179 | ↓ 19.30 | 35.22 |
| Right Pallidum to ctx lh lateralorbitofrontal | 10.577 ± 15.272 | 5.913 ± 7.893 | ↓ 44.10 | 10.9 |
| Right Thalamus Proper to ctx lh lingual | 147.678 ± 234.185 | 55.926 ± 145.136 | ↓ 62.13 | 10.08 |
| Left Amygdala to ctx lh medialorbitofrontal | 58.035 ± 72.781 | 35.318 ± 49.586 | ↓ 39.14 | 15.75 |
| ctx lh lateralorbitofrontal to ctx lh medialorbitofrontal | 254.843 ± 228.918 | 210.917 ± 137.770 | ↓ 17.24 | 10.15 |
| Right VentralDC to ctx lh rostralanteriorcingulate | 32.860 ± 43.207 | 13.872 ± 20.805 | ↓ 57.78 | 14.48 |
| ctx lh pericalcarine to ctx lh rostralanteriorcingulate | 11.128 ± 18.902 | 4.278 ± 4.815 | ↓ 61.56 | 10.29 |
| Right Hippocampus to ctx lh temporalpole | 9.394 ± 14.762 | 4.978 ± 5.698 | ↓ 47.01 | 13.77 |
| Right Thalamus Proper to ctx rh fusiform | 469.410 ± 924.148 | 143.730 ± 394.421 | ↓ 69.38 | 15.44 |
| Right Amygdala to ctx rh fusiform | 186.894 ± 256.936 | 95.337 ± 97.007 | ↓ 48.99 | 12.43 |
| Right Thalamus Proper to ctx rh lateraloccipital | 150.482 ± 267.111 | 76.258 ± 156.107 | ↓ 49.32 | 10.25 |
| ctx lh medialorbitofrontal to ctx rh lateralorbitofrontal | 54.752 ± 90.227 | 33.474 ± 45.807 | ↓ 38.86 | 14.33 |
| Right Amygdala to ctx rh lingual | 77.005 ± 100.337 | 38.411 ± 53.468 | ↓ 50.12 | 14 |
| Right Putamen to ctx rh medialorbitofrontal | 90.504 ± 98.331 | 65.072 ± 74.933 | ↓ 28.10 | 13.5 |
| ctx lh medialorbitofrontal to ctx rh medialorbitofrontal | 341.676 ± 389.227 | 244.650 ± 229.210 | ↓ 28.40 | 15.77 |
| Left Cerebellum Cortex to ctx rh parahippocampal | 28.409 ± 42.594 | 13.991 ± 21.812 | ↓ 50.75 | 10.74 |
| Right Thalamus Proper to ctx rh parahippocampal | 161.666 ± 258.747 | 60.472 ± 99.785 | ↓ 62.59 | 11.29 |

| Suprathreshold connections | (i) Connectivity strength (mean ± SD) HC, N = 509 | BD, N = 450 | (ii) Percentage change in BD (%) vs. controls | (iii) The magnitude of network component effect (t-value) |
|---|---|---|---|---|
| Right Amygdala to ctx rh parahippocampal | 201.951 ± 294.773 | 72.791 ± 98.561 | ↓ 63.96 | 14.07 |
| Right Caudate to ctx rh posteriorcingulate | 23.351 ± 40.234 | 34.192 ± 64.589 | ↑ 46.43 | 12.52 |
| Right VentralDC to ctx rh rostralmiddlefrontal | 46.663 ± 72.616 | 38.819 ± 63.012 | ↓ 16.81 | 10.45 |
| Right Pallidum to ctx rh superiorfrontal | 215.583 ± 269.257 | 134.655 ± 158.083 | ↓ 37.54 | 12.16 |
| ctx lh paracentral to ctx rh superiorfrontal | 86.490 ± 132.191 | 44.466 ± 54.818 | ↓ 48.59 | 10.98 |
| Right Pallidum to ctx rh insula | 319.502 ± 411.575 | 123.045 ± 176.904 | ↓ 61.49 | 10.52 |
| Right Accumbens area to ctx rh insula | 130.845 ± 146.007 | 70.767 ± 86.161 | ↓ 45.92 | 11.63 |

**Table S10. Number of streamlines-weighted subnetwork graph component showing dysconnectivity in bipolar disorder relative to controls.** Table shows the set of connections comprised in the subnetwork graph component found to show a significant effect for the main effect of diagnosis. NBS 'F-test' covarying for age and gender showing a dysconnected component in the bipolar group relative to controls ($t$>10, $p_{FWE}$=.005, *d=.5*). (i) Connectivity strength NOS for both groups and relative (ii) percentage decrease in strength for BD compared with controls; (iii) the magnitude of subnetwork component difference (t-value). This network showed 6.88–69.38% reduced connection strength associated with NOS-edge weighting, and 12.44-54.56% increased connection strength in BD versus controls; components of the network ranged in magnitude of difference from a T>10.08 to 35.22, with the highest effects within basal ganglia connections. BD, bipolar disorder; HC, healthy controls; NBS, network-based statistics; SD, standard deviation; ventral DC, ventral diencephalon.; ctx, cortical.

| Clinical variable | Metric | Statistics | Network type |
|---|---|---|---|
| | **Whole-brain Analysis** | | |
| **Longer illness duration** | Lower Density | β=-.1, $p_{FDR}$=.008 | Unweighted |
| | Lower Efficiency | β=-.1, $p_{FDR}$=.014 | Unweighted |
| | | β=-.1, $p_{FDR}$=.008 | NOS-weighted |
| | Longer Path length | β=-.2, $p_{FDR}=6\times10^{-37}$ | Unweighted |
| | | β=.1, $p_{FDR}$=.002 | NOS-weighted |
| **Later illness onset** | Longer Path length | β=-.1, $p_{FDR}=1\times10^{-17}$ | Unweighted |
| | | β=.1, $p_{FDR}=6\times10^{-5}$ | NOS-weighted |
| | Lower Efficiency | β=-.1, $p_{FDR}=4\times10^{-5}$ | NOS-weighted |
| **History of psychosis** | Lower Density | d=-.3, $p_{FDR}$=.0002 | Unweighted |
| | Lower Efficiency | d=-.2, $p_{FDR}$=.0002 | Unweighted |
| | | d=-.5, $p_{FDR}$=.01 | NOS-weighted |
| | Longer Path length | d=.2, $p_{FDR}$=.001 | Unweighted |
| | Higher Betweenness centrality | d=.2, $p_{FDR}$=.0002 | Unweighted |
| | **Subnetwork Analysis** | | |
| **Longer illness duration** | ↓ connectivity (right hippocampus to right amygdala) | β=-.1, $p_{FDR}$=.004 | FA-weighted |
| | ↓ connectivity (right cerebellum to right thalamus) | β=-.2, $p_{FDR}$=.02 | NOS-weighted |
| **Later illness onset** | ↓ connectivity (right cerebellum to right thalamus) | β=-.2, $p_{FDR}$=.01 | NOS-weighted |
| | ↓ connectivity (right amygdala to right accumbens) | β=-.1, $p_{FDR}$=.03 | NOS-weighted |
| | ↓ connectivity (right pallidum to right insula) | β=-.2, $p_{FDR}$=.01 | NOS-weighted |
| | ↓ connectivity (right amygdala to right fusiform) | β=-.2, $p_{FDR}$=.01 | NOS-weighted |
| | ↓ connectivity (left amygdala to left medial orbitofrontal cortex) | β=-.2, $p_{FDR}$=.01 | NOS-weighted |
| **Number of manic episodes** | ↑ connectivity (right rostralmiddlefrontal cortex to amygdala) | β=-.5, $p_{FDR}$=.001 | NOS-weighted |
| | **Medications** | | |
| **Antidepressants** | Lower Density | d=-.3, $p_{FDR}$=.036 | Unweighted |
| | Longer Path length | d=.2, $p_{FDR}$=.036 | Unweighted |
| | Lower Efficiency | d=-.3, $p_{FDR}$=.036 | Unweighted |
| | Higher Betweenness centrality | d=.2, $p_{FDR}$=.036 | Unweighted |
| **Anticonvulsants** | ↑ connectivity (right pallidum to right ventral diencephalon) | d=.4, $p_{FDR}$=.003 | NOS-weighted |
| | ↑ connectivity (right pallidum to superiorfrontal cortex) | d=.02, $p_{FDR}$=.005 | NOS-weighted |
| | ↓ connectivity (right putamen to right medial orbitofrontal cortex) | d=-.04, $p_{FDR}$=.003 | NOS-weighted |
| | ↓ connectivity (right thalamus to right fusiform) | d=.1, $p_{FDR}$=.01 | NOS-weighted |
| | ↓ connectivity (right thalamus to lingual cortex) | d=-.01, $p_{FDR}$=.01 | NOS-weighted |
| | ↓ connectivity (right thalamus to lateral occipital cortex) | d=.03, $p_{FDR}$=.01 | NOS-weighted |
| **5HT-reuptake inhibitors (NbN4)** | ↓ Density | d=-.5, $p_{FDR}$=.0003 | Unweighted |
| | ↓ connectivity (right thalamus to right hippocampus) | d=.1, $p_{FDR}$=.01 | FA-weighted |
| **DOPA/5HT/NA antagonists (NbN2)** | ↓ connectivity (left to right middle orbitofrontal gyrus) | d=.03, $p_{FDR}$=.01 | NOS-weighted |

**Table S11. Clinical associations and whole-brain/subnetwork connectivity measures in the BD group.** β=standardized regression coefficients, *d*=Cohen's *d*. NOS=number of streamlines weighted network, FA=Fractional Anisotropy-weighted network.
